# Model-driven design and evolution of non-trivial synthetic syntrophic pairs

**DOI:** 10.1101/327270

**Authors:** Colton J. Lloyd, Zachary A. King, Troy E. Sandberg, Ying Hefner, Connor A. Olson, Patrick V. Phaneuf, Edward J. O’brien, Adam M. Feist

**Affiliations:** Department of Bioengineering, University of California, San Diego, 9500 Gilman Drive, La Jolla, CA 92093, USA; Novo Nordisk Foundation Center for Biosustainability, Technical University of Denmark, 2800 Lyngby, Denmark; Bioinformatics and Systems Biology Program, University of California, San Diego, 9500 Gilman Drive, La Jolla, CA 92093, USA

## Abstract

Synthetic microbial communities are attractive for applied biotechnology and healthcare applications through their ability to efficiently partition complex metabolic functions. By pairing auxotrophic mutants in co-culture, nascent *E. coli* communities can be established where strain pairs are metabolically coupled. Intuitive synthetic communities have been demonstrated, but the full space of cross-feeding metabolites has yet to be explored. A novel algorithm, OptAux, was constructed to design 66 multi-knockout *E. coli* auxotrophic strains that require significant metabolite cross-feeding when paired in co-culture. Three OptAux predicted auxotrophic strains were co-cultured with an L-histidine auxotroph and validated via adaptive laboratory evolution (ALE). Time-course sequencing revealed the genetic changes employed by each strain to achieve higher community fitness and provided insights on mechanisms for sharing and adapting to the syntrophic niche. A community model of metabolism and gene expression was utilized to predict the relative community composition and fundamental characteristics of the evolved communities. This work presents a novel computational method to elucidate metabolic changes that empower community formation and thus guide the optimization of co-cultures for a desired application.

## Author Summary

Understanding the fundamental characteristics of microbial communities has far reaching implications for human health and applied biotechnology. Currently, many basic characteristics underlying the establishment of cooperative growth in bacterial communities have not been studied in detail. The presented work sought to elucidate the properties of nascent community formation by first employing a novel computational method to generate a comprehensive catalog of *E. coli* mutants that require a high amount of metabolic cooperation to grow in community. Three mutants from this catalog were co-cultured with a proven auxotrophic partner *in vivo* and evolved via adaptive laboratory evolution. In order to successfully grow, each strain in co-culture had to evolve under a pressure to secrete a metabolite required by the partner strain, as well as evolve to effectively utilize the required metabolite produced by its partner. The genomes of the successfully growing communities were sequenced, thus providing new insights into the genetic changes accompanying the formation and optimization of the new communities. A computational model was further developed to predict how fundamental protein constraints on cell metabolism could impact the structure of the community, such as the relative abundances of each community member.

## Introduction

Microbial communities are capable of accomplishing many intricate biological feats, due to their ability to partition metabolic functions among community members. For this reason, studying their characteristics has far reaching implications. For example, these microbial consortia have the attractive potential to efficiently accomplish complex tasks that a single engineered microbial strain likely could not. Past applications include applying communities to aid in waste decomposition, fuel cell development, and the creation of biosensors [1]. In the field of metabolic engineering, microbial communities have now been engineered that are capable of enhancing product yield or improving process stability by partitioning catalytic functions among community members [2–8]. Beyond biotechnology applications, studying microbial communities also has important health implications. This includes providing a better understanding of the gut microbiome and how it is affected by diet and other factors [9,10]. For example, metabolic cross-feeding in communities has been shown to have a role in modulating the efficacy of antibiotics treatments [11]. Developing new computational and experimental methods to understand the dynamics of microbial community formation and the inherent characteristics of established communities could therefore have far reaching implications.

Previous efforts have been made to construct synthetic communities and study their interactions and new metabolic capabilities. One such study encouraged synthetic symbiosis between *E. coli* strains by co-culturing an L-isoleucine auxotroph with a L-leucine auxotroph [12,13]. In doing so, it was found that the community was able to grow in glucose minimal media without amino acid supplementation due to amino acid cross-feeding between the mutant pairs. Mee *et al.* expanded upon this work by studying all possible binary pairs of 14 amino acid auxotrophs and developing methods to predict the results of combining the auxotrophic strains into 3-member, 13-member, and 14-member communities [14]. On a larger scale, Wintermute *et al.*co-cultured 46 conditionally lethal Keio collection *E. coli* single knockouts [15]. This effectively demonstrated that synthetic mutualism was possible in strains beyond amino acid auxotrophs [16]. These studies effectively demonstrate that new communities can be established in a relatively short time (<4 days) by pairing auxotrophic strains.

In addition to demonstrating that synthetic communities can be established, nascent auxotrophic communities can be optimized by adaptive laboratory evolution (ALE) [17]. Expanding upon the experimental work done in Mee *et al.* [14], Zhang *et al.* performed ALE on one of the co-culture pairs: a L-lysine auxotroph paired with a L-leucine auxotroph [17]. Separate co-cultures evolved to growth rates 3-fold greater than the parent, which was accomplished by forming different auxotroph strain abundances within the community. These results may have implications on both the set of metabolites being cross-fed as well as the magnitude of metabolite secretion/uptake. The increase in the evolved community growth rate is encouraging from a metabolic engineering point of view because it suggests that these binary systems can be optimized via ALE. Presumably, as an effect of an evolution, the rate of secretion/uptake of the cross-fed metabolite must increase as well to achieve higher community growth. Co-culture pairs composed of different microbial species have also been evolved with similar results [18]. Community optimization by ALE, however, has not been performed on co-cultures designed to require higher fluxes of metabolic cross-feeding than the direct biomass requirement of a particular amino acid, and the genetic mechanisms that lead to improved community fitness have not been assessed.

Established computational methods to study microbial communities often make use of genome-scale metabolic models (M-models) [19,20]. Computational models have been created and simulated using multicompartmental flux balance analysis (FBA) [21–23], dynamic flux balance analysis (dFBA) [17,24], dFBA integrated with spatial diffusion of extracellular metabolites (COMETS) [25], and FBA with game theory [26]. Novel algorithms have also been developed to describe community dynamics, such as OptCom [27], which employs a bilevel linear programming problem by maximizing community biomass as well as maximizing the inner biomass objectives of each individual species [28]. Additional ecological models have been formulated to describe community dynamics [29–31]. Despite the advances made by these approaches, the role of efficient proteome allocations in driving community dynamics has not been studied in detail.

Here, we demonstrate that nascent *E. coli* communities can be constructed from co-cultures of auxotroph mutants requiring high fluxes of metabolic cross-feeding. We first introduce the OptAux algorithm for designing auxotrophic strains that require high amounts of supplemented metabolites in order to grow (**Figure 1A**). The OptAux solutions provided a catalog of starting strains from which four auxotrophic mutants were selected to co-culture and optimized via adaptive laboratory evolution (**Figure 1B**). In optimizing the growth of the nascent co-culture communities, significant metabolic rewiring had to occur to allow the strains to cross-feed the high levels of the necessary metabolites. The genetic changes accompanying this rewiring was assessed by analyzing the genetic changes (mutations and observed genome region duplications). This analysis thus enabled predictions of primary metabolite cross-feeding and community composition.

**Figure 1.**
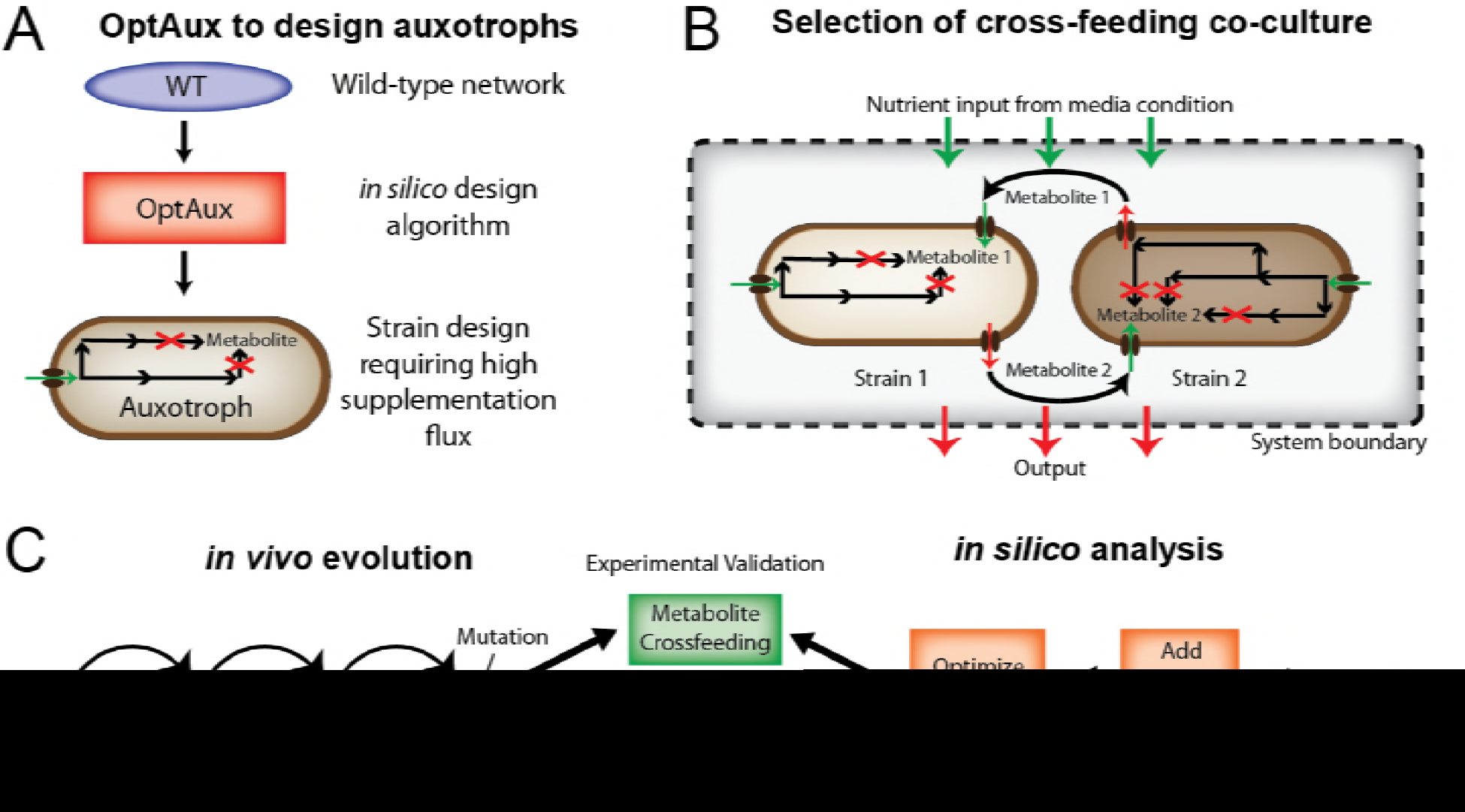
Study Overview. **A**: An algorithm was developed to *de novo* predict reaction deletions that will produce *E. coli* strains auxotrophic for a metabolite of interest. **B**: From the set of auxotrophic strain designs, pairs were selected to determine whether they were capable of forming a nascent syntrophic cross-feeding community. **C**: The chosen co-cultures were both evolved via adaptive laboratory evolution and modeled using a genome-scale model of *E. coli* metabolism and expression (ME-model) [19,20]. The model predictions of optimal strain abundances and metabolite cross-feeding were verified using resequencing data from the co-culture wet-lab experiments.

To study the characteristics of designed and optimized communities, a community model of metabolism and expression (ME-model) was constructed [32–34] (**Figure 1C**). Such a modeling approach was necessary since previous methods of genome-scale community modeling have focused on studying the metabolic flux throughout community members (using M-models) without consideration of the enzymatic cost of proteins and pathways that drive these metabolic processes. As proteome optimization via niche partitioning and cell specialization is a driving factor of community formation in ecological systems [35–38], it is essential to consider proteomic constraints when studying bacterial communities. To this end, community ME-models were successfully utilized to interpret the nascent communities and were used to suggest approaches to optimize the evolved co-cultures and potentially modulate metabolic crossfeeding.

## Results

### OptAux Development and Simulation

The OptAux algorithm identifies strain designs that are predicted to be auxotrophic for a metabolite of interest. The algorithm was built by modifying an existing concept introduced for the design of metabolite producing strains [39] which was later additionally implemented in a mixed-integer linear programming (MILP) algorithm (RobustKnock [40]). Three key modifications were made to derive OptAux from RobustKnock. **First**, the inner growth rate optimization was replaced so that OptAux can be run at a predetermined minimum growth rate bound (*set_biomass* constraint **Figure 2B**). This ensures that OptAux designs are auxotrophic at all growth rates (**Figure 2A**). **Second**, the objective coefficient was reversed in order to allow the algorithm to optimize for metabolite uptake as opposed to secretion. **Third**, a constraint was added to allow the model to uptake any additional metabolite that can be consumed by the model (*trace_metabolite_threshold* constraint **Figure 2B**). For simulations in which this threshold value was set above zero, all possible exchange metabolites included in the model had their lower bound set to the *trace_metabolite_threshold* value to compete with a target metabolite uptake, allowing the “specificity” of the knockout solution to Be adjusted. Specificity, in this case, refers to whether the mutant strain will be auxotrophic for a given metabolite in the presence of other metabolites. High specificity solutions are auxotrophic for only one metabolite, regardless of whether other metabolites are present. With this implementation, OptAux identified strain designs that require the targeted metabolite at all growth rates with varying metabolite specificity and uptake requirements.

**Figure 2.**
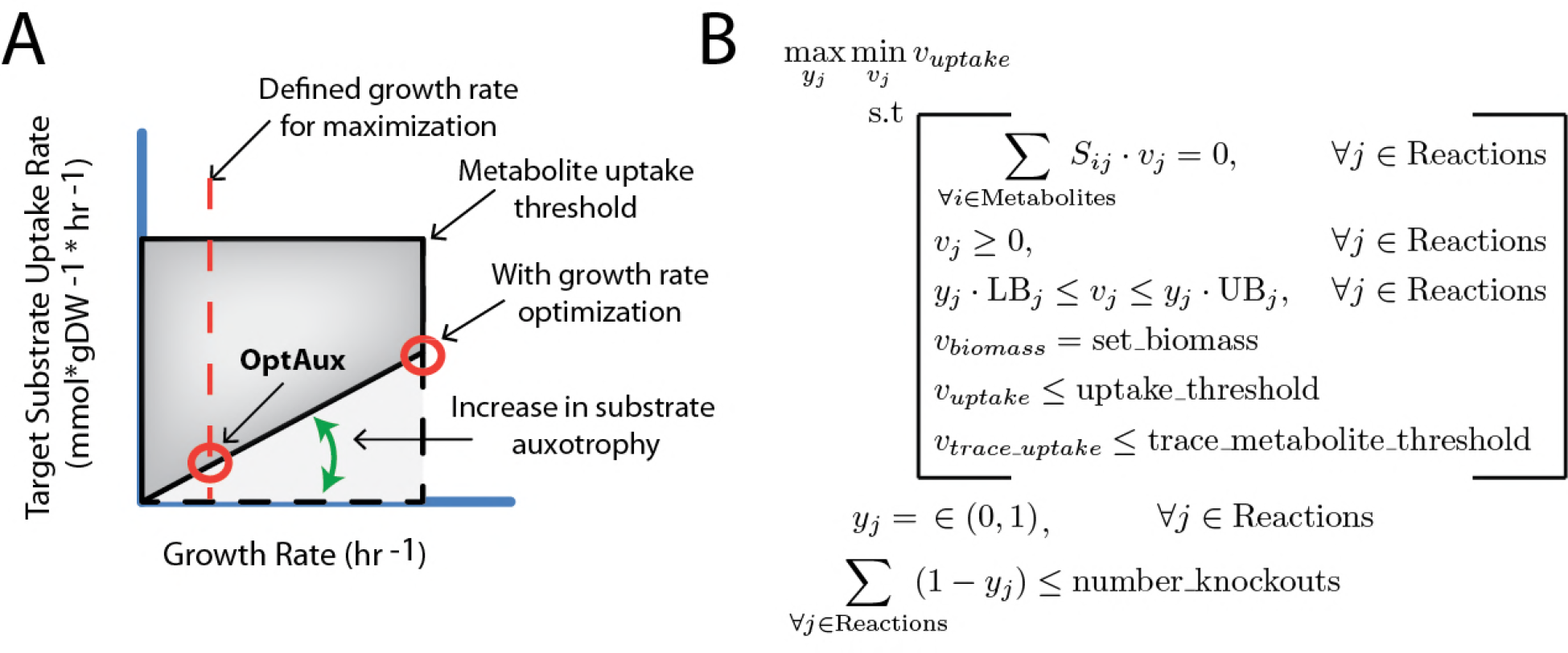
OptAux Design. **A**: OptAux was developed to maximize the minimum uptake of a target metabolite. Unlike algorithms such as OptKnock with tilting [21] and RobustKnock [22], this optimization occurs at a predetermined growth rate as opposed to using an inner optimization of growth rate (depicted with the red circles). This is to ensure that all OptAux designs will be auxotrophic for the target metabolite at all growth rates, particularly low growth rates. The dotted lines show the required uptake for the metabolite with no genetic interventions. In this case, uptake of the target metabolite is not required at any growth rate. The solid black lines depict the maximum and minimum uptake required for a particular metabolite of an OptAux designed strain. **B**: The OptAux optimization problem. For further description of the algorithm and underlying logic see **Methods**.

OptAux was utilized on the *i*JO1366 M-model of *E. coli* K-12 MG1655 [41] to comprehensively examine auxotrophic strain designs. OptAux was run with 1, 2, and 3 reaction knockouts for 285 metabolite uptake reactions using 4 different trace metabolite thresholds (**S1 Data**). Of the given solutions, 228 knockout sets were found to be capable of producing 66 unique strain auxotrophies. This set of strain designs presents an expansive look into the auxotrophies possible in the *E. coli* K-12 MG1655 metabolic network, which could be used to understand the possible niches of *E. coli* could inhabit in natural or synthetic communities [42].

### OptAux Solution Characteristics

The OptAux strain designs were broken into two major categories based on the number of metabolites which, when supplemented, restore cell growth: **1) Essential Biomass Component Elimination Designs (EBC**, **Figure 3B)** and **2**) **Major Subsystem Elimination Designs (MSE**, **Figure 3A**). The EBC designs are characterized as auxotrophic strains with high metabolite specificity. They were broken into two subcategories: specific auxotrophs (only one metabolite can restore growth, **Figure S2**) which consists of 104 (23 unique) knockout sets, and nonspecific auxotrophs (defined as strains in which less than 5 metabolites can restore growth, **Figure S2**) which consists of 55 (20 unique) knockout sets. The specific and nonspecific EBC designs were preferred at high trace metabolite threshold values. There is significant overlap between OptAux predicted EbC designs, and known *E. coli* auxotrophic mutants [14,43–54]. A summary of experimentally characterized OptAux designs are presented in **Table S1**. Of note, there are five designs that were not found to be previously characterized in the scientific literature, and these present novel *E. coli* auxotrophs.

**Figure 3.**
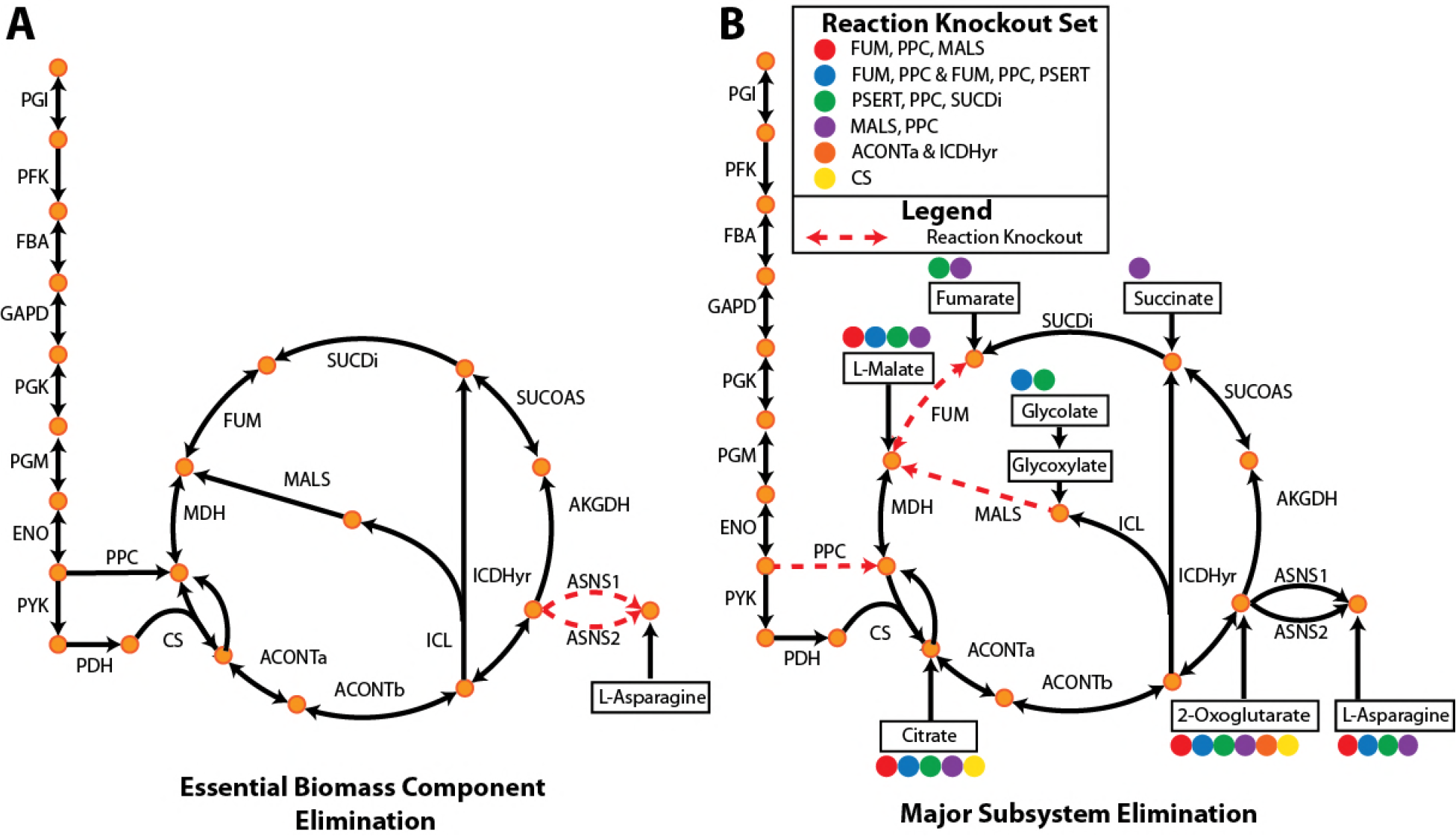
OptAux Solutions. Depending on the parameters used when running OptAux, two major solution types are possible. **A: Essential Biomass Component Elimination** designs, like the ASNS1 and ASNS2 knockout shown, can grow only when one specific metabolite is supplemented. For the case shown, this metabolite is L-asparagine. **B**: Alternatively, **Major Subsystem Elimination** designs have a set of alternative metabolites that can restore growth in these strains. Examples of these designs are shown for citric acid cycle knockouts sets. One specific three reaction knockout design (FUM, PPC, MALS) is shown in red dashed lines where four metabolites in the figure can individually rescue this auxotroph (marked with solid red circles). The metabolites that can restore growth for each of the knockout strain designs listed in the legend are indicated with the colored circle associated with the reaction knockouts.

MSE designs were analyzed as novel auxotrophic strain designs. These were defined as strains in which five or more metabolites could restore growth and consisted of the remaining 69 (23 unique) sets of knockouts. At low trace metabolite thresholds, MSE designs were the preferred OptAux solution. This knockout strategy was often accomplished through knockouts to block metabolic entry points into anabolic subsystems. One such example of an MSE design is given in **Figure 3B**. Here a three reaction knockout design of the FUM, PPC, and MALS reactions can be rescued by one of the four compounds in the figure (citrate, L-malate, 2-oxoglutarate, or L-asparagine) at an average required uptake flux of 0.4 mmol gDW^−1^ hr^−1^ to grow at a rate of 0.1 hr^−1^. These rates are higher than the fluxes needed to rescue the EbC design in **Figure 3A**, which requires uptake of 0.024 mmol gDW^−1^ hr^−1^ on average to grow at a rate of 0.1 hr^−1^. Another design was a glutamate synthase (GLUSy) and glutamate dehydrogenase (GLUDy) double knockout which effectively blocks the entry of nitrogen into amino acid biosynthesis by preventing its incorporation into 2-oxoglutarate to produce L-glutamate. This renders the cell unable to produce all amino acids, nucleotides, and several cofactors. In order to grow at a rate of 0.1 hr^−1^, this strain is computationally predicted to require one of 19 individual metabolites at an average uptake of 0.62 mmol/gDW/hr (**Supplemental Data File 2**).

MSE designs are of particular interest as they are largely unique, nontrivial, and have often not been studied as *E. coli* auxotrophies. However, some of the MSE single knockouts have been used for large-scale studies of auxotrophic co-culture short term growth [16]. Since these predicted MSE knockouts disrupt significant biological processes, they produce auxotrophies that require much larger amounts of metabolite supplementation in order to grow, compared to EBC designs (e.g., **Figure S3**). This makes MSE *E. coli* mutants attractive from a microbial community perspective because they would require a pronounced rewiring of the metabolic flux of their partner stains in co-culture to secrete the high amount of the auxotrophic metabolite needed for community growth.

### Adaptive Laboratory Evolution of Auxotrophic *E. coli* Co-cultures

To demonstrate how the OptAux algorithm can be leveraged to design strains and co-culture communities, *E. coli* auxotrophic mutants were validated in the wet lab and evolved in coculture. Three communities were tested, each consisting of pairwise combinations of four OptAux predicted auxotrophs. This included one EBC design, Δ*hisD*, which was validated as an L-histidine auxotroph, paired with each of three MSE designs, Δ*pyrC*, Δ*gltA*Δ*prpC*, and Δ*gdhA*Δ*gltB*. These three MSE strains had disruptions in pyrimidine synthesis, TCA cycle activity, and nitrogen assimilation into amino acids, respectively (**Table S2**). The Δ*pyrC* mutant was computationally predicted to be able to grow when supplemented with one of 20 metabolites in *i*JO1366, and the Δ*gltA*Δ*prpC* and Δ*gdhA*Δ*gltB* mutants were predicted to grow in the presence of 14 and 19 metabolites, respectively (**S2 Data, Table S4**).

Upon inoculation into the first flask of batch growth, each of the co-culture’s growth rates were low (<0.05 hr^−1^) suggesting the strains initially showed minimal cooperativity or metabolic crossfeeding (**Figure S4**). Following approximately 40 days of ALE, all three co-culture combinations had evolved to establish a nascent community, indicated by an increase in the co-culture growth rate. There was diversity in the endpoint batch growth rates among the independently evolved triplicates for each of the Δ*hisD* & Δ*pyrC* and the Δ*hisD* & Δ*gdhA*Δ*gltB* co-cultures with endpoint growth rates ranging from 0.09-0.15 hr^−1^ and 0.08-0.15 hr^−1^, respectively. The four successfully evolved independent replicates for the Δ*hisD* & Δ*gltA*Δ*prpC* co-cultures also showed endpoint growth rate diversity ranging from 0.12−0.19 hr^−1^ (**Table 1**, **Figure 4A**). The relatively large range in endpoint growth rates for all co-cultures suggests that a subset of replicates evolved to a less optimal state and could be further improved if given more time to evolve.

**Figure 4.**
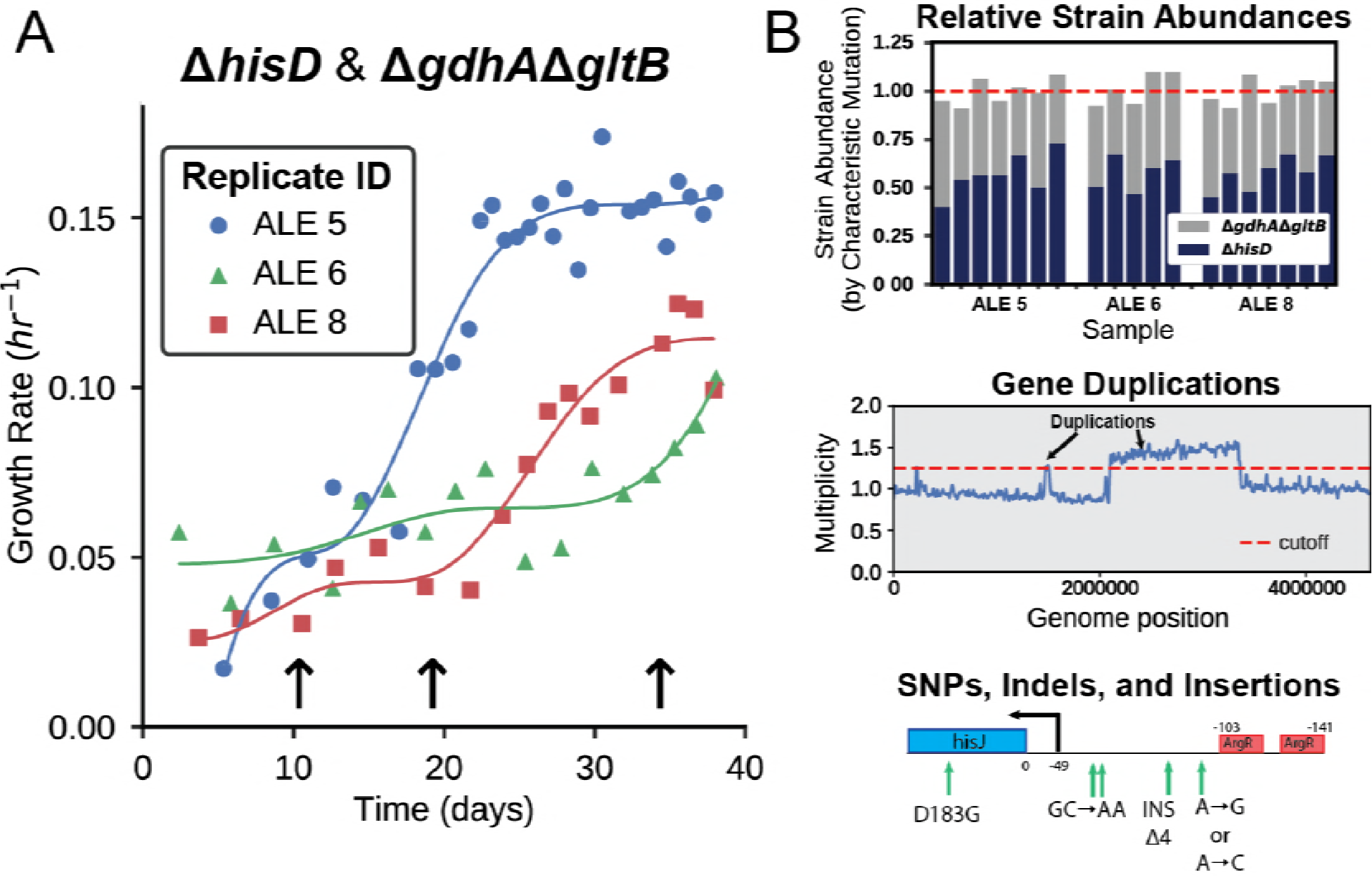
Adaptive Laboratory Evolution. **A**: *E. coli* co-cultures were evolved over a 40 day period and the growth rate was periodically measured. Three of the co-cultures evolved were capable of establishing syntrophy and showed an improvement in growth rate. The arrows indicate the time points at which samples were taken during the Δ*hisD* & Δ*gdhA*Δ*gltb* co-culture ALE. **B**: Each of the sampled co-cultures were sequenced. This information was used to predict the fractional strain abundances of each of the co-culture members. It was also used to identify duplications in genome regions of one of the community members and infer causal mutations that improved community fitness.

**Table 1.**
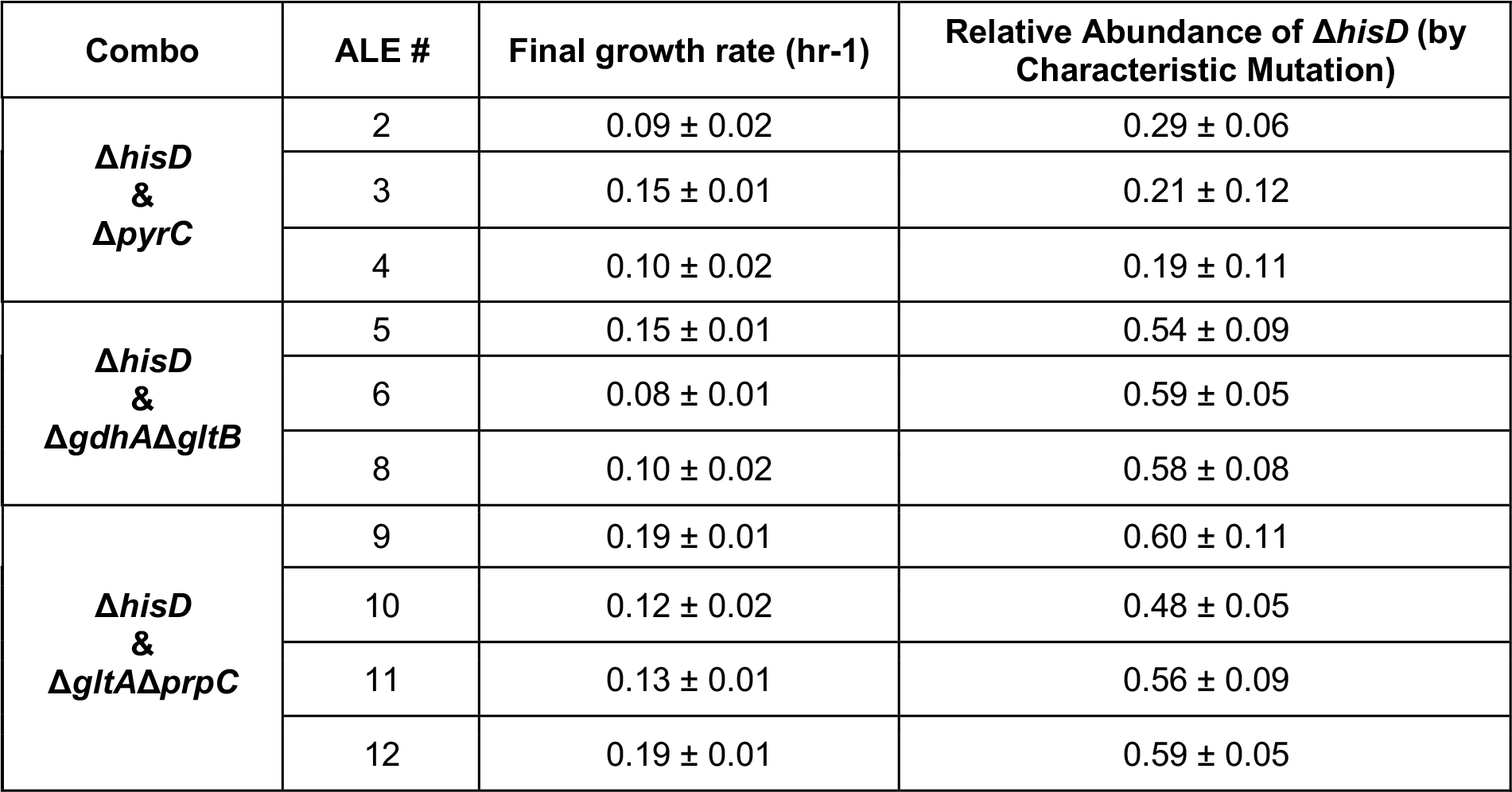
Final growth rates and fractional strain abundance of the Δ*hisD* strain, by characteristic mutation, for each ALE lineage.

| Combo | ALE # | Final growth rate (hr<sup>-1</sup>) | Relative Abundance of $\Delta hisD$ (by Characteristic Mutation) |
| --- | --- | --- | --- |
| **$\Delta hisD$**<br>**&**<br>**$\Delta pyrC$** | 2 | $0.09 \pm 0.02$ | $0.29 \pm 0.06$ |
| | 3 | $0.15 \pm 0.01$ | $0.21 \pm 0.12$ |
| | 4 | $0.10 \pm 0.02$ | $0.19 \pm 0.11$ |
| **$\Delta hisD$**<br>**&**<br>**$\Delta gdhA\Delta gltB$** | 5 | $0.15 \pm 0.01$ | $0.54 \pm 0.09$ |
| | 6 | $0.08 \pm 0.01$ | $0.59 \pm 0.05$ |
| | 8 | $0.10 \pm 0.02$ | $0.58 \pm 0.08$ |
| **$\Delta hisD$**<br>**&**<br>**$\Delta gltA\Delta prpC$** | 9 | $0.19 \pm 0.01$ | $0.60 \pm 0.11$ |
| | 10 | $0.12 \pm 0.02$ | $0.48 \pm 0.05$ |
| | 11 | $0.13 \pm 0.01$ | $0.56 \pm 0.09$ |
| | 12 | $0.19 \pm 0.01$ | $0.59 \pm 0.05$ |

To probe the metabolic strategies of the three co-culture pairs, the genomes of the populations were resequenced at several time points over the course of the 40 day evolution (**Figure 4A**). The resequencing data was used to identify gene region duplications and acquired mutations (**Figure 4B**) that provided insight into the specific mechanisms employed by the co-cultures to establish cooperation.

The relative strain abundance of each mutant was also tracked to understand the dynamics of community composition in the synthetic co-culture. Each starting strain contained unique characteristic mutations (**Table S3**) which could act as a barcode to track the community composition (**Figure 4b**, **Table 1**). The breseq mutation identification software [55] was used to calculate the frequency of each of these characteristic mutations within a sequenced co-culture. The relative frequency of the characteristic mutations was used to approximate the fraction of each strain within the co-culture population. This analysis showed that 2 of the 3 co-culture combinations maintained similar relative fractions of the two member strains, whereas one coculture, Δ*hisD* & Δ*pyrC*, consistently maintained a relative Δ*pyrC* abundance of near three quarters of the total population (71-81%, **Table 1**). Alternatively, the relative abundance of each strain in the populations was predicted by comparing the read coverage of the deleted genes relative to the mean, which showed good agreement with the characteristic mutation-based predictions (**Figures S4-5**).

### Mutations Likely Affecting Metabolite Uptake/Secretion

Several evolutionary strategies were observed in the mutations identified across the ten successfully evolved co-culture lineages (**Tables S5-7**). One ubiquitous strategy across all three co-culture pairs was to acquire mutations within or upstream of inner membrane transporter genes. For instance, numerous mutations were observed in every co-culture lineage in the *hisJ* ORF or in the 5’UTR of the *hisJ* operon. This operon contains all four genes (*hisJ, hisM, hisP, hisQ*) composing the histidine ABC uptake complex, the primary mechanism for histidine uptake in *E. coli* K-12 MG1655 [56]. Seven mutations were found in the region directly upstream of the transcription start site (**Figure 5**). Three of the five substitutions were observed in more than one co-culture pairing with a SNP in one position (A->G, A->C or A->T at 86 base pairs upstream of *hisJ*) appearing to be particularly beneficial as it was identified in every lineage except one (ALE #5). In three ALEs, a mutation was observed within the *hisJ* ORF that resulted in a substitution of aspartate residue at the 183 position by glycine. Based on the protein structure, this substitution could disrupt two hydrogen bond interactions with bound L-histidine ligand in the periplasm [57]. Further mutations were observed that could affect the binding of the ArgR repressor to the 5’ UTR of the *hisJMPQ* operon or affect the activity of the ArgR protein itself (**Table S5**). This included a 121 base pair deletion and a SNP in the binding site of the ArgR repressor in the 5` UTR of *hisJ* (**Figure 5**). The mutation in the *argR* ORF consisted of a frameshift insertion early in the coding sequence and persisted throughout ALE #8, appearing in the Δ*hisD* endpoint clone (**Table S6**). ArgR functions to repress L-arginine uptake and biosynthesis as well as the L-histidine ABC uptake complex [58] in response to elevated L-arginine concentrations. All of these mutations could improve L-histidine uptake in the Δ*hisD* strains either by directly increasing the efficacy of the HisJMPQ ABC uptake system or by preventing ArgR mediated repression of this transporter.

**Figure 5.**
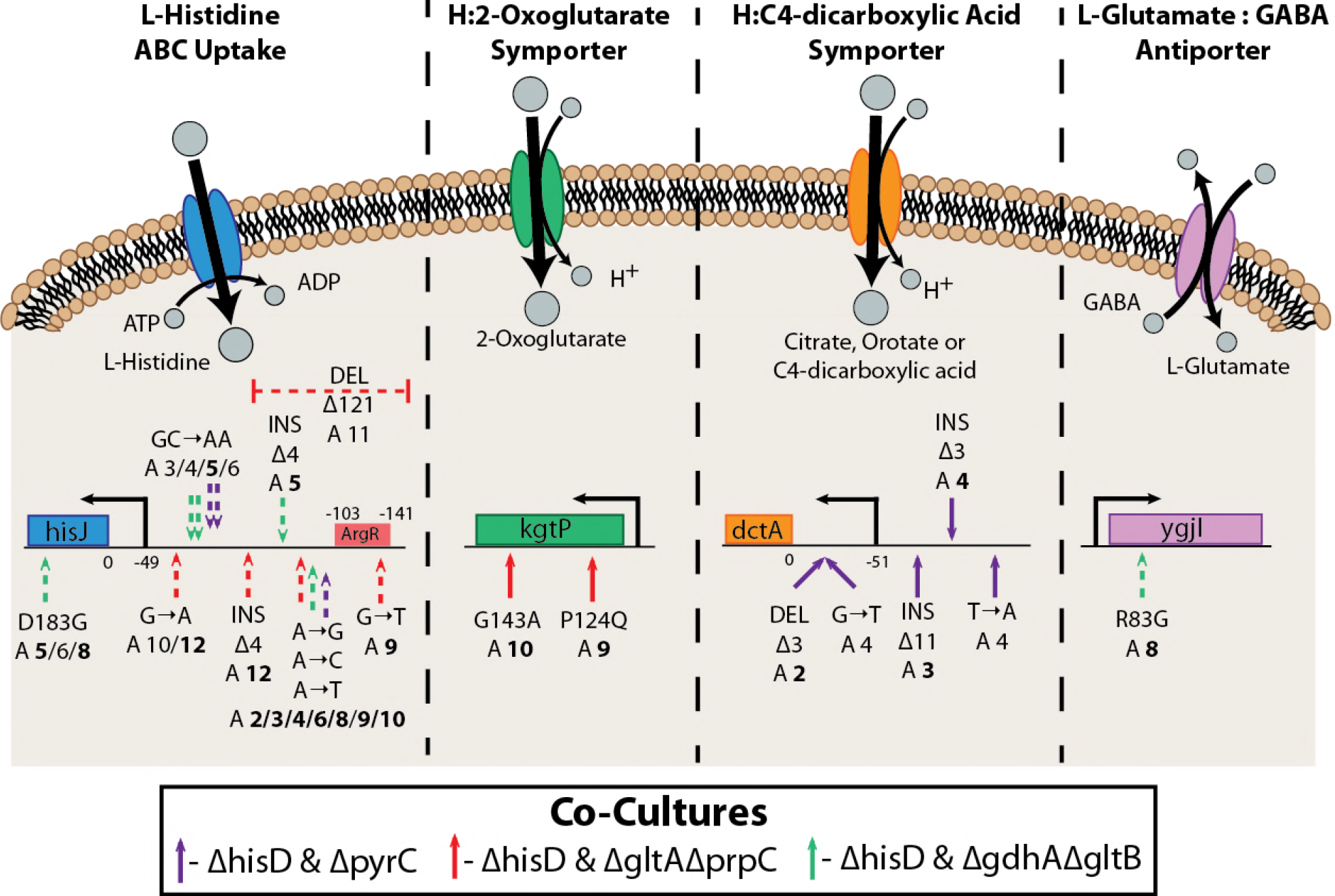
Mutations Affecting Inner Membrane Metabolite Transport. Mutations were observed affecting the activity of four inner membrane transporters. A schematic of the function or putative function of each transporter is shown. Depicted below the schematics are the relative locations of the observed mutations on the operon encoding each of the enzymatic complexes. For example, all ten evolved Δ*hisD* strain endpoints possessed at least one mutation in or upstream of *hisJ*. This operon includes genes coding for HisJMPQ, the four subunits of a histidine ABC uptake system. A depiction of the activity of this complex is shown, in which energy from ATP hydrolysis is used to transport histidine into the cytosol from the periplasm. Mutations are indicated on the operon schematics if mutations appear at >10% frequency in more than one flask, and ALE numbers are in bold if the mutation appears in the endpoint clone strain. The mutations indicated with a dashed arrow occurred in the Δ*hisD* strain and a solid arrow indicates they occurred in its partner MSE strain.

Beyond improving the uptake of L-histidine in the Δ*hisD* strain, mutations were observed that could improve metabolite uptake in a partnering strain. For instance, in the Δ*hisD* & Δ*gltA*Δ*prpC* co-culture, two of the Δ*gltA*Δ*prpC* strains acquired mutations in the *kgtP* ORF (a transporter of 2-oxoglutarate [59]) that were present in the endpoint clones. These mutations include a substitution of a L-proline residue with a L-glutamine at the 124 position and a substitution of a glycine residue with an L-alanine at the 143 position. These two substitutions occurred in the fourth transmembrane helix in the protein and a cytoplasmic region [60], respectively, and could act to augment or complement the mutation in the 5’ UTR of the *kgtP* ORF seen in the starting clone of the Δ*gltA*Δ*prpC* mutant (**Table S5**). Both the accumulation of mutations associated with this transporter and the fact that the citrate synthase knockout mutant is computationally predicted to grow in the presence of 2-oxoglutarate suggest that Δ*gltA*Δ*prpC* could be cross-fed 2-oxoglutarate when in co-culture.

A recurrent mutation was observed in the Δ*hisD* & Δ*pyrC* co-culture that could function to better facilitate uptake of a metabolite being cross-fed from the Δ*hisD* strain to the Δ*pyrC* strain. The three independently evolved lineages each acquired at least one mutation in the 5’ UTR of *dctA*, which were confirmed to be in all Δ*pyrC* endpoint clones (**Table S7**). The gene product of *dctA* functions as a proton symporter that can uptake orotate, malate, citrate, and C4-dicarboxylic acids [61] (**Figure 5**). Further, simulations of a Δ*pyrC* strain predict that growth is possible with orotate supplementation, but not with any of the other metabolites known to be transported by the *dctA* gene product. Thus, it was proposed that these mutations could act to increase the activity of this transporter to allow the Δ*pyrC* strain to more efficiently uptake orotate cross-fed by the Δ*hisD* strain.

Lastly, one lineage of the Δ*hisD* & Δ*gdhA*Δ*gltB* co-culture acquired a SNP in the *ygjI* coding region; the SNP was present in the Δ*hisD* endpoint clone and resulted in a substitution of L-arginine for glycine at position 83 of this protein. This position is within a periplasmic region and one residue prior to a transmembrane helix of the protein [62]. The function of this protein has not been experimentally confirmed, but based on sequence similarity, it is predicted to be a GABA:glutamate antiporter [63]. Given that this mutation was seen in the Δ*hisD* clone, it is possible that this mutation had the effect of increasing the strain’s secretion of 4-aminobutyrate (GABA) or L-glutamate. Such a mutation could improve community fitness by facilitating the cross-feeding of either these metabolites to the Δ*gdhA*Δ*gltB* strain since it is predicted to be auxotrophic for both metabolites (**Table S4**).

### Mutations Likely Affecting Nitrogen Regulation

Removing reactions in major biosynthetic pathways likely results in a disruption of the homeostatic concentrations of key sensor metabolites or an activation of nutrient limitation stress responses. Mutations were observed in the evolved co-cultured sets which point to mechanisms to adapt to these pathway disruptions. Examples of this adaptation included three frameshift mutations early in the *glnK* ORF found in three Δ*gltA*Δ*prpC* clones from the Δ*hisD* & Δ*gltA*Δ*prpC* co-cultures (**Figure 6A**) and one premature stop codon in the Δ*hisD* clone of the same co-culture. The frameshift mutations would possibly affect the AmtB nitrogen uptake system as well, as it is located on the operon downstream of GlnK. GlnK is one of two nitrogen regulators with overlapping functions that are uridylated depending on the relative concentrations of 2-oxoglutarate and L-glutamate. In conditions of high relative 2-oxoglutarate concentration relative to L-glutamate, GlnK is uridylated and the *E. coli* nitrogen limitation response is triggered [64]. Unlike the alternative nitrogen regulator, GlnB, however, when not uridylated GlnK binds to the Amtb nitrogen uptake complex, reducing its activity [65]. The citrate synthase knockout (*Δ*gltA*Δ*prpC**) in particular could see a disruption in the homeostatic concentrations of metabolites immediately downstream of the reaction, including 2-oxoglutarate and L-glutamate. This could impair the ability of the cell to respond to sensors of nitrogen excess or limitation and respond with the proper global regulatory changes. Removing the activity of this GlnK mediated response system would prevent any detrimental cellular responses given that the strains are both grown in excess ammonium. No mutations, however, were observed in the alternative nitrogen regulator, GlnB, throughout any of the evolutions.

**Figure 6.**
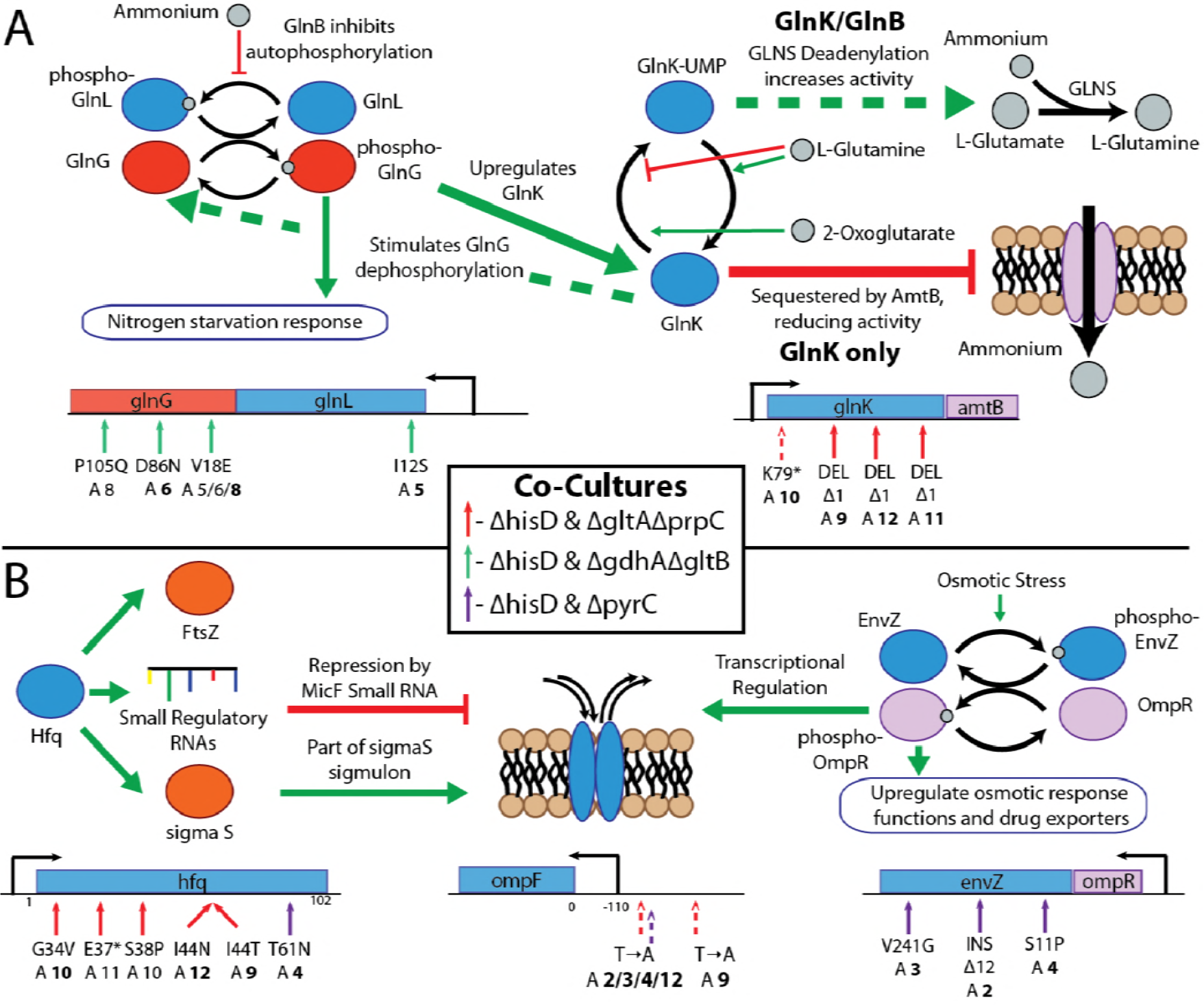
Mutations Affecting Stress Responses and Metabolite Homeostasis. Functions or putative functions of mutated genes are summarized with schematics with the location of all mutations on the operon below the schematic. Mutations are indicated on the operons schematic if mutations appear at >10% frequency in more than one flask and, ALE numbers are in bold if the mutation appears in the endpoint clone strain. The mutations indicated with a dashed arrow occurred in the Δ*hisD* strain and solid arrow if they occurred in its partner MSE strain. **A**: Mutations observed related to nitrogen starvation and metabolite homeostasis. Mutations were acquired within the open reading frame of both genes comprising the nitrogen sensing two-component regulatory system. Shown in the schematic is the regulatory cascade in which nitrogen is sensed by GlnL, which stimulates its autophosphorylation and subsequent donation of the phosphorus group to GlnG. Phosphorylated GlnG upregulates general functions associated with nitrogen starvation. Further, mutations were observed in the ORF of GlnK, one of two nitrogen regulators, sharing most functions with Glnb. Both genes become uridylylated in response to high concentrations of 2-oxoglutarate and low concentrations of glutamine, which is an indication of low nitrogen concentration. GlnK-UMP can activate GLNS deadenylation, thus increasing its activity. Unlike Glnb, GlnK when in a deuridylylated state (high concentrations of glutamine) can be sequestered by the Amtb ammonium transporter causing it to have a reduction in activity [26] Dashed lines in the schematic indicate primary GlnB functions and solid lines indicate primary GlnK function. **B**: Mutations observed associated with the *E. coli* stress response. Numerous mutations were observed in the ORF of Hfq which is an RNA-binding protein with numerous global functions. These include interactions with small regulatory RNAs which are often required to enable the small RNA’s regulatory function. Hfq is also required for the wild-type expression of the S sigma factor. Both MicF and sigma S are involved in regulating the expression of outer membrane porin *ompF*, a gene which acquired mutations in multiple ALE lineages. Mutations were also observed in the *envZ* ORF which is the sensory protein in the osmotic stress two-component regulatory system. Upon sensing osmotic stress, it autophosphorylates and transfers a phosphate to OmpR, thus upregulating osmotic stress genes. These genes consist of many outer membrane porins, including ompF.

Mutations found in the Δ*gdhA*Δ*gltb* strains imply a change in the activity of the two-component nitrogen regulatory system. This strain in all Δ*hisD* & Δ*gdhA*Δ*gltb* lineages acquired mutations in the open reading frame of at least one gene in the two-component nitrogen regulator system, glnG (ntrC) and glnL (ntrb) (**Figure 6A**) [64]. Amino acid substitutions were observed in position 18, 86, and 105 of *glnG* corresponding to the response receiver domain of GlnG, likely augmenting its ability interact with GlnL (based on protein families [66]). The endpoint clone of ALE #5 acquired an amino acid substitution of L-isoleucine to L-serine within a PAS domain of GlnL at position 12. This corresponds to the protein domain where regulatory ligands likely bind [67] so this mutation could act to augment its autophosphorylation activity in response to nitrogen. Given the location of these mutations, it can be hypothesized that they functioned to decrease the regulatory activity of the two-component system response to excess nitrogen. For the Δ*gdhA*Δ*gltB* strain, this could be beneficial to reduce the GlnGL mediated downregulation of nitrogen uptake and assimilation processes.

Mutations were also observed affecting osmotic as well as nonspecific stress responses (**Figure 6b**). These are summarized in the **Supplemental Text**.

### Genome Duplications Complement Sequence Changes

A complementary adaptive strategy for improving co-culture community fitness was to acquire duplications in regions of the genome (**Figures S7-9**). In some cases, this evolutionary strategy appeared to function to amplify expression of transporters (also observed in [68]) to more efficiently uptake a metabolite that can rescue the strain’s auxotrophy. Alternatively, these duplications could function to increase the likelihood of acquiring mutations in the duplicated region [69,70]. Therefore, the genes contained within the duplicated regions in some cases provided clues to which metabolites were cross-fed within the co-culture. For example, one of the three Δ*hisD* & Δ*gdhA*Δ*gltB* lineages displayed clear increases in coverage near positions 674-683 kbp and 1,391-1,402 kbp with multiplicities exceeding 15. The former of these coverage peaks included 9 genes, including the 4 genes composing the GltIJKL L-glutamate/L-aspartate AbC uptake system [71]. The latter peak included 10 genes including the 4 genes in the *abgRABT* operon, which facilitates the uptake and hydrolysis of p-aminobenzoyl-glutamate into glutamate and 4-aminobenzoate [72]. This suggests that both of these metabolites could be cross-fed to the Δ*gdhA*Δ*gltB* strain, though the *abgRABT* duplication was depleted in favor of the *gltIJKL* duplication over the course of the evolution, suggesting L-glutamate or L-aspartate is the preferred cross-feeding metabolite over p-aminobenzoyl-glutamate (**Figure 7**).

**Figure 7.**
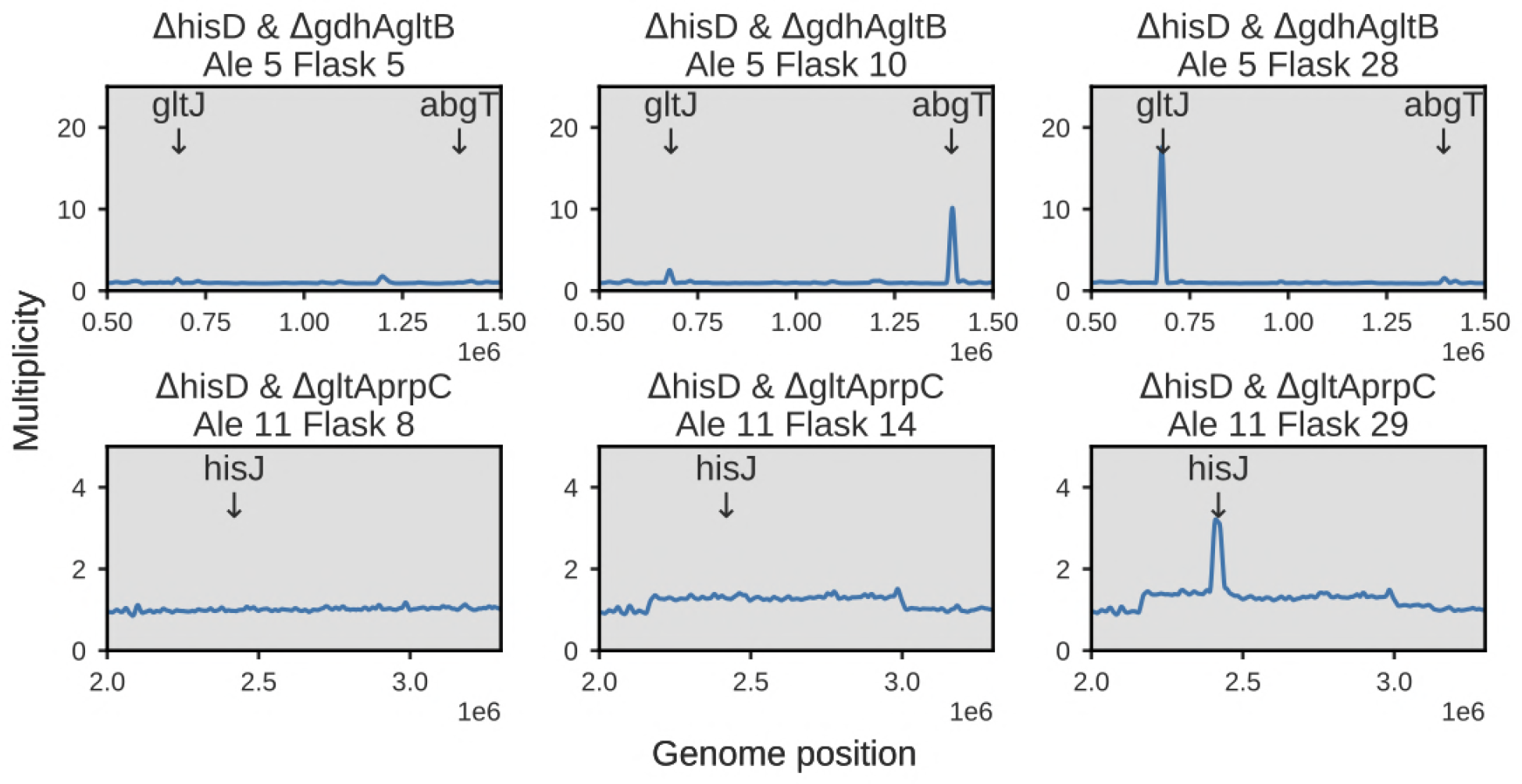
Duplication Dynamics. The top panel depicts the dynamics of two high multiplicity duplications in two transport complexes in *E. coli* throughout the course of Ale #5 of a Δ*hisD* & Δ*gdhA*Δ*gltB* pair. A small region containing the *abgT* symporter of p-aminobenzoyl glutamate is duplicated early in the evolution, but is later replaced by duplications in a region containing *gltJ* and the rest of the genes comprising the GltIKJL L-glutamate/aspartate AbC uptake system. The bottom panel depicts the course of Ale #11, a Δ*hisD* & Δ*gltA*Δ*prpC* co-culture, which initially showed a broad ~1Mbp duplication. By the end of the evolution either a nested duplication emerged or a significant subpopulation emerged that contained a duplication of a small genome region containing *hisJ* and the rest of the HisJMPQ L-histidine ABC uptake system.

While the duplications mentioned above presented clear amplifications in targeted operons, some observed duplications consisted of 100,000s of basepairs and 100s of genes. Further, many of the duplications seen in the populations were not observed in the sequenced endpoint clones. Possible explanations for these observations can be found in the **Supplemental Text**.

### Modeling Community Features of Auxotroph Communities

Community ME-models were created for each of the three evolved co-culture sets (**Figure S10**). The models were constructed based on the assumption that, in order to form a stable community when growing exponentially, the strains in co-culture must be growing, on average, at an equal rate. Mass balance conversion terms could then be used to relate the metabolic flux that a strain contributes to the shared compartment and its fractional abundance (see **Methods**). This approach offered a means to understand which factors drive the structure of the newly established communities (i.e., the relative abundance of the community members) and, ultimately, how this relates to metabolite cross-feeding.

The community ME-models have the capability of assessing how the community composition could vary depending on the identity of the metabolite that is cross-fed or the enzyme efficiency of the community members. The role of the cross-fed metabolites in defining the structure of the community was assessed using the community ME-models by: 1) allowing metabolic crossfeeding to remain unrestricted and 2) restricting the cross-feeding to only one metabolite. When the metabolite cross-feeding was left unrestricted (i.e., any metabolite restoring growth in either strain was allowed to cross-feed in the simulation, **Supplemental Text, Figure S11**) computed cross-feeding profiles were complex and prediction of the identity of the cross-fed metabolite did not strongly point to one potential metabolite (**Figure S12**). However, when turning to the sequencing data, there was general agreement between predicted and experimentally inferred optimal community structure which provided confidence in using the proposed modeling approach (**Figure S11**).

Alternatively, the second approach to assess the influence of metabolite cross-feeding on community composition involved restricting the simulation to cross-feed only one of the metabolites computationally predicted to restore growth in the MSE strain. In doing so, the identity of the metabolite being cross-fed could be related to the optimal community growth rate and structure. This approach additionally offered a way to narrow the set of optimal or near optimal cross-feeding metabolites that would be predicted to be cross-fed *in vivo*. The computations predicted that the Δ*hisD* & Δ*pyrC* co-culture would have a community composition and growth rate robust to the metabolite being cross-fed with a slightly higher community growth rate if orotate, uracil, uridine monophosphate, or uridine were cross-fed. The optimal composition of the community was predicted to be skewed toward low percentages (~20%) of the Δ*hisD* strain for all metabolites in this co-culture. The Δ*hisD* & Δ*gltA*Δ*prpC* and Δ*hisD* & Δ*gdhA*Δ*gltB* co-cultures, on the other hand, were sensitive to the cross-feeding metabolite where the community structure depended on the identity of the cross-feeding metabolite (**Figure 8A**). For these two co-cultures, the Δ*hisD* & Δ*gltA*Δ*prpC* and Δ*hisD* & Δ*gdhA*Δ*gltB* pairs were computationally predicted to achieve higher community growth rates when cross-feeding L-glutamate, 2-oxoglutarate, citrate, or L-glutamine and 4-aminobutanoate, L-aspartate, L-glutamine, L-glutamate, L-alanine, or L-asparagine, respectively.

**Figure 8.**
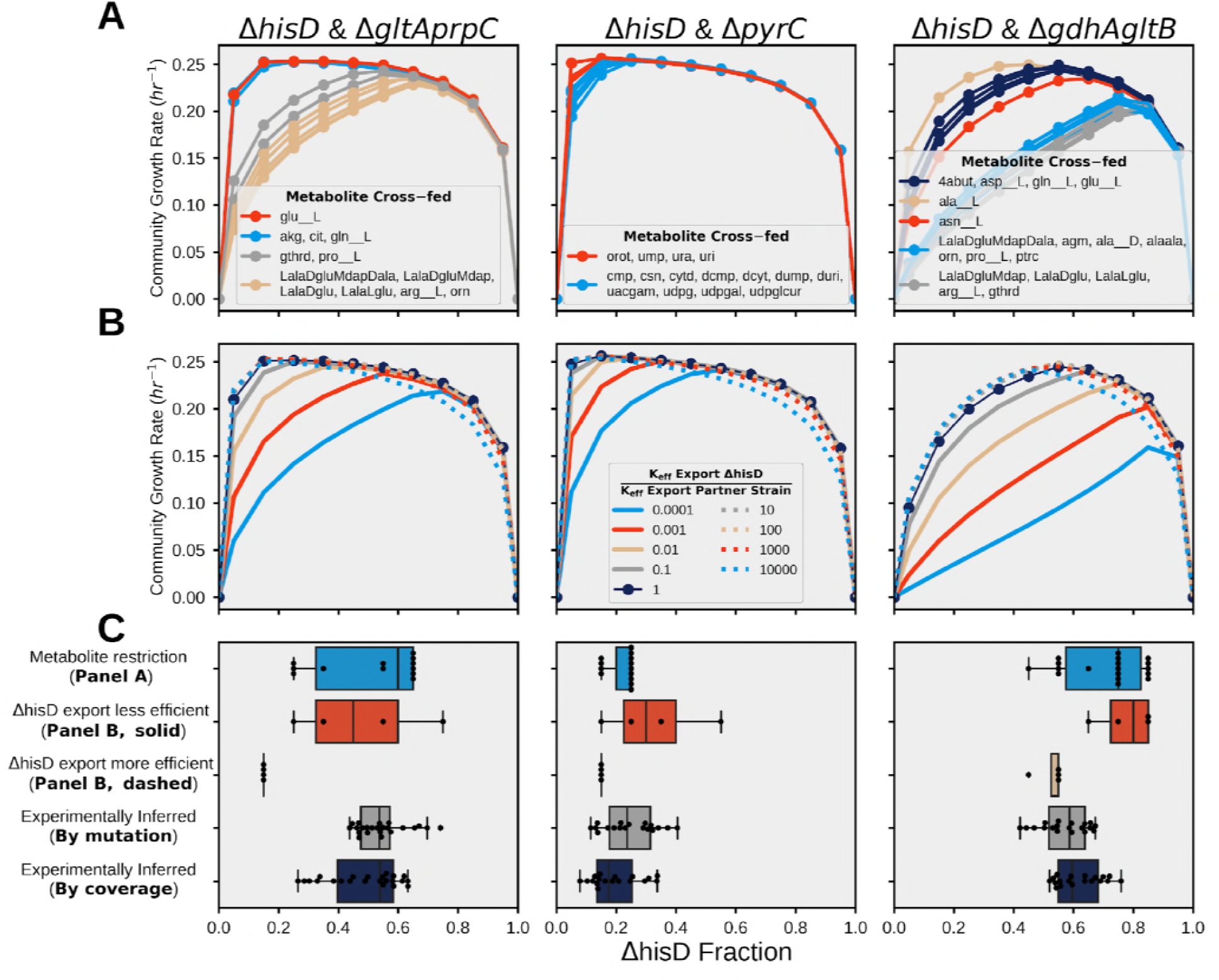
Community Modeling. Community ME-model predicted growth rates for fractional strain abundances of Δ*hisD* ranging from 0 to 1. **A**) The effect of metabolite cross-feeding on community structure. Each curve was computed by allowing different metabolites to be cross-fed to the MSE strain. Similar curves were grouped by color. **B**) Effect of varying the proteome efficiency of metabolite export on community structure (see **Methods**). The analysis was performed on models constrained to only cross-feed the metabolite that was inferred from the resequencing data (2-oxoglutarate, orotate, and L-glutamate, respectively) (**Table 2**). **C**) Box plots of experimentally measured abundances for each sample (bottom two rows, gray, and dark blue) and the computationally-predicted optimal strain abundances following variation in the cross-feeding metabolite (top row, blue) and in strain proteome efficiency (second and third row, red, and yellow).

Community ME-models further enable an examination of how each strain’s proteome “efficiency” may affect co-culture characteristics when growing in its community niche. Such an analysis was performed by altering a ME-model parameter for each strain corresponding to how efficiently it can export the metabolite that is cross-feeding its partner strain (see **Methods**). This parameter can be used as a proxy for cellular proteome investment in wasteful or inefficient processes when synthesizing and exporting a metabolite, which is likely to occur in substantial amounts until the strains further adapt to grow as a community. That is, the cells will not be able to optimally rearrange their proteome and metabolic fluxes to efficiently grow as a community over this short-term evolution. It is possible, however, that some strains in co-culture will be able to reorganize their proteome to secrete the necessary metabolite more or less efficiently than their partner strain (**Table 2**). The proteome efficiency analysis showed that the community compositions of all three co-cultures were moderately sensitive to this parameter (**Figure 8B**). Further, the pairs showed a bimodal behavior depending on whether the Δ*hisD* strain was more or less efficient than its partner (**Figure 8B**). The community models predicted that if the export processes of the Δ*hisD* strain require a greater protein investment relative to the default export efficiency parameter, the abundances of the Δ*hisD* strain will increase in the community. Conversely, if the partner strain requires greater protein investment, the community composition remains stable and unchanged. The optimal predicted community composition for the two analyses shown in Figure 8A **and** B are summarized in **Figure 8C**. The figure shows general agreement between the computed optimal community compositions and the experimentally inferred community composition, even after varying key features of the community simulation. This suggests that community ME-models have the potential to be useful tools for understanding the behavior of simple communities.

**Table 2.** Metabolite being cross-fed by the Δ*hisD* strain to its partner strain, as inferred from sequencing data.

| Pair with $\Delta hisD$ | Inferred Metabolite | Mutation Evidence | Duplication Evidence |
| --- | --- | --- | --- |
| $\Delta pyrC$ | Orotate | Mutations in 5' UTR of *dctA* in $\Delta pyrC$ strain in all ALEs (**Figure 5**) | Broad duplication in portion of genome containing *dctA* coding region in all ALEs (**Figure S9, S4 Data**) |
| $\Delta gdhA\Delta gltA$ | L-Glutamate | Ale #8 mutation in *ygjI* ORF in $\Delta hisD$ strain (**Figure 5**) | ALE #5/6 targeted duplications in *gltJ* coding region (**Figure 7, Figure S8**)<br>ALE #5 transient duplication in *abgT* coding region (**Figure 7**) |
| $\Delta gltA\Delta prpC$ | 2-Oxoglutarate | Starting mutation in 5' UTR of *kgtP* in $\Delta gltA\Delta prpC$ strain (**Table S5**)<br>ALE #9/10 Acquired mutations in *kgtP* ORF in $\Delta gltA\Delta prpC$ strain (**Figure 5**) | - |

## Discussion

This study has demonstrated a novel workflow to design, optimize, and computationally interpret non-trivial syntrophic co-cultures to better understand the characteristics of simple microbial community formation. The simple communities consisted of two strains of *E. coli* K-12 MG1655 which required, in order to grow themselves, the growth of their partner strain. To design the communities to possess characteristics more attractive from an engineering perspective, a novel algorithm, termed OptAux, was used. This algorithm was used to design highly auxotrophic strains which, when paired in co-culture, require high levels of metabolic crossfeeding in order for the community to grow. Three co-cultures consisting of OptAux designs were tested *in vivo* and optimized via adaptive laboratory evolution. By analyzing the genetic changes observed throughout the evolution we could infer the cellular changes underlying improvements in the fitness of the highly metabolically-coupled communities. This work thus provided new insight into cellular mechanisms for establishing syntrophic growth. A community ME-model was developed to computationally interpret the communities and their fundamental properties. Such models are the first to offer a means to study, on the genome-scale, how efficient proteome allocation to metabolic functions in the community members can influence the structure of the nascent microbial communities.

### OptAux Can be Used to Design Novel Communities

To facilitate the design of co-culture communities requiring significant metabolic rewiring and cross-feeding, we constructed the OptAux algorithm to find reaction knockouts that will create auxotrophic strains requiring high amounts of metabolites for growth (**Figure 2**). OptAux returned two kinds of solutions depending on the parameters used, so-called Major Subsystem Elimination (MSE) and Essential biomass Component Elimination (EBC) designs (**Figure 3**). EBC designs are specific with regard to which metabolites are required for the strain to grow and correspond to auxotrophs that have been validated in previous studies [14,50–54]. OptAux EBC predictions resulted in eight designs that were previously verified experimentally and five predictions of untested auxotrophs (**Table S1**). Conversely, the MSE designs are computationally predicted to grow when supplemented with a any of variety of different metabolites and represent largely new designs that have not been characterized experimentally, though some of the single gene knockout MSE designs were grown in co-culture in [16]. MSE auxotrophs in co-culture need high levels of cross-feeding in order to grow (0.05 and 0.2 mmol gDW^−1^ hr^−1^ on average for an EBC and MSE strain to grow at a rate of 0.1 hr^−1^, respectively), requiring significant metabolic rewiring in its partner strain (**Figure S13**).

### ALE was Successfully Applied to Increase Fitness of Co-culture

Four OptAux predicted auxotrophic *E. coli* mutants were constructed *in vivo*, confirmed as auxotrophs, and grown in co-culture. A growth rate selection pressure was applied on these nascent, poorly growing communities via ALE. Three co-cultures of an Δ*hisD* EBC strain paired with an MSE strain showed reproducible improvements in growth rate throughout the course of the ALE (**Table 1**). Under these conditions each of the strains had to rewire its metabolic network to both secrete a metabolite required by its partner strain and efficiently import the metabolite needed to grow itself through mutations that were identified, effectively establishing a new microbial community. By selecting for growth rate, a novel indirect selection pressure was applied on each strain to increase the secretion and uptake of the cross-fed metabolites, thus improving the growth of the co-culture community. This evolution design therefore has potential as a system to self-optimize microbial strains as industrial producers of metabolites of interest.

Throughout the course of adaptive laboratory evolution, the nascent communities improved community fitness by acquiring beneficial mutations (**Tables S5-7, Figures S7-9**). There was a high degree of parallelism in the identified shared mutations and duplications which appeared in each co-culture pair’s ALE lineages, providing confidence that the acquired mutations and duplications were meaningful and causal in improving community fitness [73]. Consistently, duplications coincided with genome regions containing mutations acquired in endpoint clones. It has been shown that, as a mechanism for evolving new cellular functions, microbes duplicate genome regions to provide the redundancy needed for divergence of function or for acquiring new or altered capabilities [70]. Further, similar gene duplications in nutrient transporters have been shown in yeast to provide fitness benefits in glucose limited environments by increasing the expression of the transporter [68].

Beyond enabling an analysis of how the co-cultures were capable of establishing syntrophy, the sequencing data provided a measure of the structure of the community in terms of relative strain abundance. All auxotrophic mutants contained a unique characteristic starting mutation (**Table S3**), which was used to track the relative abundance of each member of the co-culture community throughout the evolutions. Community structures appeared to remain remarkably consistent both across ALE replicates of the same strain combinations and over time throughout the ALE lineages (**Table 1**, **Figures S4-5**). This finding was corroborated by using the coverage of the gene deletion regions in population resequencing (**Figures S5-6**). The observation of stable community composition is in line with what has been observed in multi-species microbial soil communities grown in single substrate minimal media [74].

### Resequencing Data Provides Insight into Probable Metabolite Cross-feeding

Mutational evidence, often related to transporter processes, from the evolved populations provided insight into which metabolites were being cross-fed within the co-cultures. For instance, all ALE lineages acquired mutations targeting the ABC uptake system for L-histidine (**Figure 5**). Given that all of the three evolved co-culture sets included a strain that was an EBC auxotroph for L-histidine, community growth logically would increase if histidine uptake was improved in this strain via these genetic changes. Similarly, the three MSE strains that were paired with the L-histidine auxotroph, Δ*pyrC*, Δ*gdhA*Δ*gltB* and Δ*gltA*Δ*prpC*, displayed evidence in their resequencing data to suggest that the strains were being cross-fed orotate, glutamate and 2-oxoglutarate, respectively (**Table 2**). A community ME-model was constructed for each of the three communities and the model simulations predicted a hierarchy, where clusters of metabolites provide slight benefits in predicted community growth rates relative to other metabolites. In each case, the mutation data inferred cross-feeding metabolites were contained in one of the top computationally predicted clusters.

### Community ME-modelling Allows for Analyzing Co-culture Composition

Community ME-models were employed to understand how the proteome efficiency of each strain drives community composition. ME-models are uniquely capable of addressing this question because they directly incorporate the proteomic cost of catalyzing a metabolic process, which is particularly necessary in this system as there is an inherent proteome cost of each strain to cross-feed the necessary metabolite in co-culture [75]. Kinetic parameters, which play a role in dictating proteome cost in these community ME-models, were therefore systematically adjusted to understand how each strain’s proteomic “efficiency” affected the simulation characteristics. The simulations predicted that, for all of the three co-cultures, the proteomic efficiency of the Δ*hisD* would have the largest impact on the relative abundance of each coculture member (**Figure 8B**). This is an expected finding due to the fact that the Δ*hisD* strain has the larger cross-feeding burden since it is paired with an MSE strain in each case. Further, when the Δ*hisD* secretion proteome efficiency was decreased, the community ME-model predicted its optimal abundance in the co-culture would actually increase. Though unintuitive, this prediction is in agreement with a paradox predicted in a previous computational study of community dynamics [76]. In addition to proteome efficiency, the ME-model predicted that the identity of the metabolite being cross-fed has an effect on optimal community composition (**Figure 8A**). The distributions of possible community compositions based on varying metabolites and proteome efficiency aligned well with two of the three co-cultures (Δ*hisD* & Δ*gdhA*Δ*gltB* and Δ*hisD* & Δ*pyrC*, **Figure 8C**). This implies that community ME-modelling potentially offers a means to study how changes in the characteristics of each strain in coculture will affect the optimal community structure and growth behavior.

From an industrial perspective, shifting the community composition could increase the production of a specific metabolite of interest. Therefore, this modeling method offers a way to predict how, for instance, LacZ or other unused (i.e., non beneficial) proteins could be efficiently overexpressed to lower a strain’s proteome efficiency and alter community composition, thus improving the yield of metabolite secretion. Additionally, this modeling method suggests that the identity of the cross-feeding metabolite can bias the optimal community composition to some extent. For instance, for the Δ*hisD* & Δ*gdhA*Δ*gltB* co-culture, the Δ*hisD* relative fraction can vary from 0.45 to 0.85 if L-alanine is cross-fed versus L-arginine. By somehow biasing the crossfeeding toward one metabolite or the other (e.g., exporter knockout), the community composition could potentially be manipulated, thus altering the yield of the cross-fed metabolite.

## Conclusions

This work demonstrated a novel approach using both a design algorithm and community modeling to understand how strains adapt to grow in new community niches. The work also provided insight into evolutionary strategies bacteria can use to readjust their metabolism and respond to drastic changes in homeostatic metabolite concentrations while learning to inhabit this new biological niche. Beyond better understanding ecological communities, this workflow could be applied as a tool for developing new platform bacterial strains for producing metabolites of industrial relevance. Lastly, the novel community resource allocation model was successfully used to predict co-culture community characteristics. This modeling tool could be leveraged to predict experimental strategies for optimizing a community to fit the desired application and have broad impacts on human health [77,78].

## Materials and Methods

### Computational Methods

All constraints based modeling analyses were performed in Python using the COBRApy software package [79] and the *i*JO1366 metabolic model of E. coli K-12 MG1655 [41]. For aerobic simulations the maximum oxygen uptake rate was constrained to 20 mmol • dDW^−1^ • hr^−1^, and the maximum substrate was constrained to 10 mmol • dDW^−1^ • hr^−1^. All *i*JO1366 optimizations and algorithm solutions presented were found using the Gurobi (Gurobi Optimization, Inc., Houston, TX) mixed-integer linear programming (MILP) or linear programming (LP) solver. The community ME-models were solved using the qMINOS solver in quad precision [87,88]. All scripts and data used to create the presented results can be found at www.github.com/coltonlloyd/optaux.

#### OptAux Algorithm Formulation

The OptAux algorithm was derived based on the ideas from existing MILP algorithms (i.e., RobustKnock [40] and OptKnock [80]). A new algorithm was written as opposed to implementing a “reverse” version of RobustKnock where the algorithm would optimize the uptake of a metabolite at the maximum growth rate. A “reverse” RobustKnock implementation would lead to strain designs that must take up a high amount of the target metabolite when approaching the maximum growth rate (**Figure S1A**). In order for a strain to be truly auxotrophic for a particular metabolite, however, it must be required at all growth rates (**Figure 2A**, **Figure S1B**). To ensure that OptAux designs have this auxotrophic phenotype, the inner problem optimizing for growth rate utilized in RobustKnock was replaced with a *set_biomass* constraint. This forced the metabolite uptake optimization to occur at a predefined growth rate and was implemented by setting the upper and lower bounds of the biomass objective function to this value:

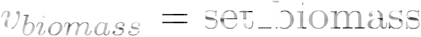

For the simulations ran in this study (**S1 Data**), the *set_biomass* value was set as 1/10 the maximum growth rate for the wild-type simulation in *in silico* glucose minimal media supplemented with the metabolite whose uptake is being maximized.

An additional constraint was applied to represent additional metabolites present in the media. It was applied by finding all metabolites with exchange reactions with its lower bound set to zero and increases the bound to the trace_metabolite_threshold, shown for exchange reaction *i* below:

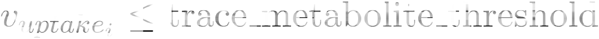

Increasing this threshold ultimately increases the specificity of the OptAux solution in regards to other metabolites that can potentially restore growth. In other words, this effectively models a scenario where, along with the presence of the target metabolite and primary substrate, there are trace amounts of competing energy source and biosynthetic precursor metabolites in the *in silico* media.

The resulting algorithm is a bilevel MILP **Figure 2B**) that can be found at www.github.com/coltonlloyd/optaux.

#### OptAux Simulations

The OptAux algorithm was ran for all carbon containing metabolites with exchange reactions in *i*JO1366. For each optimization the target metabolite is selected and the maximum uptake of the metabolite is set to 10 mmol/gDW/hr. The model was then reduced by performing flux variability analysis (FVA) on every reaction in the model and setting the upper and lower bounds of each reaction to the FVA results. If FVA computed that no flux could be carried through the reaction, then it was removed from the model. Additionally, reactions were excluded from knockout consideration if they met one of the following criteria: **1)** it is a /JO1366 false positive when glucose is the primary carbon substrate [81] **2)** it is essential in LB rich media [15] **3)** its annotated subsystem is one of the following: Cell Envelope Biosynthesis, Exchange, Inorganic Ion Transport and Metabolism, Lipopolysaccharide Biosynthesis / Recycling, Murein Biosynthesis, Murein Recycling, Transport, Inner Membrane, Transport, Outer Membrane, Transport, Outer Membrane Porin, or tRNA Charging **4)** it involves a metabolite with more than 10 carbons **5)** it is a spontaneous reaction.

#### Identifying Gene Mutations and Duplications

The FASTQ data from the samples sequencing was filtered and trimmed using AfterQC version 0.9.6 [82]. The quality controlled reads were aligned to the genome sequence of *E. coli* K-12 BW25113 (CP009273.1) [83] using Bowtie2 version 2.3.0 [84]. Mutations were identified based on the aligned reads using breseq version 0.32.0b [55]. If the sample was of a co-culture population and not a clone, the predict polymorphism option was used with a frequency cutoff of 0. 025. The output of the breseq mutation analysis for all samples can be found in **S3 Data**.

Duplications were found by analyzing the BAM sequence alignment files output from Bowtie using the pysam Python package [85]. Pysam was used to compute the sequencing read coverage at each DNA position within the genome sequence. For population samples, a cutoff of 1.25 × coverage fit mean (measure of average read alignment coverage over the genome), a relatively low threshold to account for the varying fractional abundances of the strains in community. A gene was flagged as duplicated in the sample if over 80% of the base pairs in the gene ORF had alignment coverage above the duplication threshold. Duplications found in starting strains were excluded from duplication analysis. Further the set of duplicated genes were grouped together if there are located next to each other on the genome. A new group was made if there was more than five genes separating a duplicated gene from the next duplicated gene (**S4 Data**).

Aligned contig coverage across the *E. coli* genome is noisy and therefore must be filtered before plotting in order to observe its dominant features. This was accomplished by first splitting the coverage vector into 50,000 segments such that each segment represented ~100 base pairs and the average of the segments was found. Locally weighted scatterplot smoothing (LOWESS) was then applied to the array of concatenated segments using the statsmodel package in python [86]. For the smoothing 0.5% of all of the segments was used when estimating each coverage value (y-value), and zero residual-based reweightings were performed. The remaining parameters were set to their default.

#### Calculating Strain Abundances from Resequencing Data

The fractional strain abundance of each strain in co-culture were predicted using two features of the resequencing data of each co-culture population sample: **1)** the frequency of characteristic mutations of each strain and **2)** the relative coverage of the knocked out genes.

Each of the stains used in this study possessed a unique characteristic mutation (**Table S3**), which could be used as a barcode to track the strain. The breseq population mutation calling pipeline would identify the characteristic mutations of each strain in co-culture and report the frequency that the mutation occurred. This output was used to track their presence. For strains with two characteristic mutations (Δ*hisD*, Δ*gdhA*Δ*gltB*) the average of the frequency of each gene was used as a prediction of the relative abundance of that strain. One mutation in particular, an IS element insertion in yqiC which is characteristic of the ΔhisD strain, was not detected in several samples when Δ*hisD* was in co-culture with ΔpyrC. This is likely due to the low frequency of the Δ*hisD* strain in that particular population. In those cases, the Δ*hisD* strain abundance was predicted using only the frequency of the lrhA/alaA intergenic SNP (**Figure S5**).

The second method used the contig read alignment to compare the coverage of the deleted genes in each strain to the fit mean coverage of the sample. As an example, for a strain paired with the Δ*hisD* strain, the average coverage of the base pairs in the *hisD* ORF divided by the fit mean for that sample, would give an approximation of its relative abundance in the population. As with the characteristic mutation approach, if the two genes are knocked out in the strain, the average coverage of the two genes is used to make the approximation (**Figure S5**).

When reporting the relative abundance predictions, the predicted abundances of each strain was normalized by the sum of the predicted abundances of the two strains in co-culture. This ensured that the abundance predictions summed to one. Predictions made using the two described methods showed general agreement (**Figure S6**).

#### Community Modeling

Community ME-models were created using a multicompartment FBA approach, where each of the two mutant strains in co-culture occupy a compartment with an additional shared compartment where each of the strains can exchange metabolites. The relative abundance of each strain was accounted for by adjusting the exchange reaction from a strain’s compartment into the shared compartments. For secretion, this was done by multiplying these exchange reactions as follows:

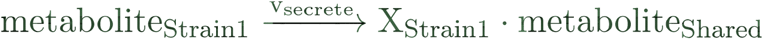

and for uptake:

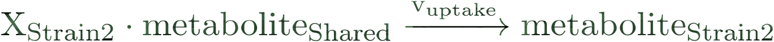

where v_secrete_ is the secretion flux from strain 1 and has units of mmol • gDW_Strain1_^−1^ • hr^−1^ and X_Strain1_ is the fractional abundance of strain 1 with units of gDW_Strain1_ • gDW_Community_. Therefore applying this coefficient to metabolite_Shared_ gives the reaction fluxes from strain 1 (v_secrete_) in units of mmol • gDW_Community_^−1^ • hr^−1^. For the subsequent uptake of the shared metabolite by strain 2, the fractional abundance of strain 2 is applied giving units of mmol • gDW_Strain2_^−1^ • hr^−1^ (**Figure S10**).

Using this community modeling approach, the fractional abundance (X_i_) of each strain in the coculture was implemented as a parameter that could be varied from 0 to 1, which in turn had on impact on the optimal growth state of the community. Simulations were ran varying X_Strain1_ (abundance of strain 1) from 0.05 to 0.95 and the community growth rate was optimized. The metabolites that were allowed to be cross-fed in simulation were limited to the set of metabolites that can computationally restore the growth of each auxotroph (**Table S4**).

For the community simulations, the *i*JL1678b [32] model of *E. coli* K-12 MG1655 was used with the uptake of metabolites in the *in silico* glucose minimal growth media into the shared compartment left unconstrained, as the ME-model is self limiting [33]. The non-growth associated ATP maintenance and the growth associated ATP maintenance were set to the default parameter values in the model. The RNA degradation constraints were removed to prevent high ATP costs at the low community growth rates. Since, the newly formed communities are highly unoptimized and growing slowly, the unmodeled/unused protein fraction parameter was set to 75%. If a metabolite had a reaction to import the metabolite across the inner membrane, but no export reaction, a reaction to transport the metabolite from the cytosol to the periplasm was added to the model. For more on the model parameters, refer to [32] and [33].

To vary the proteomic efficiency (k_eff_) of the export metabolites, first the exchange reaction into the shared compartment for all potential cross-feeding metabolites except the metabolites inferred from the experimental data (**Table 2**) was constrained to zero. Then the enzymatic efficiency of the outer membrane transport process of only the inferred metabolite was altered in each strain. The outer membrane transport reactions for each inferred metabolite (i.e.,HIStex, GLUtex, AKGtex, and OROTtex for L-histidine, L-glutamate, 2-oxoglutarate, and orotate, respectively) have multiple outer membrane porins capable of facilitating the transport process. To account for this the k_eff_ kinetic parameter of each of porin and reaction changed by multiplying the default k_eff_ value by the appropriate multiplier. The COBRAme software was used for all ME-model manipulations [32].

#### Reproducibility

All code and data necessary to reproduce the results can be found on GitHub at https://github.com/coltonlloyd/OptAux.

### Experimental Methods

#### *E. Coli* Strain Construction

All single gene knockouts used in this work were obtained from the Keio collection, a collection of all single gene knockouts in E. coli K-12 BW25113 [15]. To generate double gene knockout strains, the second knockout genes were identified from the Keio collection as donor strains, and their P1 phage lysates were generated for the transduction into the receiving single KO strains. For instance, the Δ*gltA* or Δ*gltB* knockout strain was a donor strain and the Δ*prpC* or Δ*gdhA* knockout strain was a receiving strain (**Table S2**). These four knockout strains were used for the construction of double knockout strains of Δ*gltA*Δ*prpC* and Δ*gdhA*Δ*gltB*. Each mutant was confirmed not to grow in glucose M9 minimal media without supplementation of an auxotrophic metabolite predicted by the *i*JO1366 model.

#### Adaptive Laboratory Evolution

Cultures were initially inoculated with equal numbers of cells from the two relevant auxotrophs, then serially propagated (100 μL passage volume) in 15 mL (working volume) flasks of M9 minimal medium with 4 g/L glucose, kept at 37°C and well-mixed for full aeration. An automated system passed the cultures to fresh flasks once they had reached an OD600 of 0.3 (Tecan Sunrise plate reader, equivalent to an OD600 of ~1 on a traditional spectrophotometer with a 1 cm path length), a point at which nutrients were still in excess and exponential growth had not started to taper off. Four OD600 measurements were taken from each flask, and the slope of ln(OD600) vs. time determined the culture growth rates.

#### Resequencing

Co-culture populations samples were collected at multiple points throughout the ALE and sequenced. Additionally, the starting mutant strains and both mutants isolated from the ALE endpoint samples were sequenced. The Δ*hisD* endpoint clone was unable to be isolated via colony selection for ALE #11. Genomic DNA of the co-culture populations and mutant clones was isolated using the Macherey-Nagel NucleoSpin tissue kit, following the manufacturer’s protocol for use with bacterial cells. The quality of isolated genomic DNA was assessed using Nanodrop UV absorbance ratios. DNA was quantified using the Qubit double-stranded DNA (dsDNA) high-sensitivity assay. Paired-end whole genome DNA sequencing libraries were generated using Illumina’s Kappa kit and run on an Illumina MiSeq platform with a PE600v3 kit. DNA sequencing data from this study will be made available on the Sequence Read Archive database (submission no. SUb3903910).

## Acknowledgements

We thank Richard Szubin for help preparing samples for resequencing and thank Joshua Lerman and Justin Tan for informative discussions. This research used resources of the National Energy Research Scientific Computing Center, which is supported by the Office of Science of the US Department of Energy under Contract No. DE-AC02-05CH11231. Funding for this work was provided by the Novo Nordisk Foundation through the Center for Biosustainability at the Technical University of Denmark [NNF10CC1016517]. CJL was supported by the National Science Foundation Graduate Research Fellowship under Grant no. DGE-1144086.

## Author contributions

C.J.L., Z.A.K. and A.M.F. designed the study. C.J.L. and Z.A.K. developed OptAux. C.J.L. and E.J.O. developed the community ME-modeling method. C.J.L. performed all computation and analysis. Y.H. and C.A.O. constructed all *E. coli* mutant strains and T.S. performed the adaptive laboratory evolution. C.J.L. and A.M.F. wrote the manuscript and all authors reviewed the text and provided edits.

## Conflict of interest

The authors have no conflicts of interest to declare

## Supporting Information

**S1 Data. OptAux Solutions**. Output of the OptAux algorithm ran for one, two, and three reaction knockouts on glucose minimal media for all carbon containing exchange metabolites. Four different trace metabolite thresholds were used (0, 0.01, 0.1, 2).

**S2 Data. Major Subsystem Elimination Designs**. All MSE designs along with further information regarding the subsystems of the reaction knockouts and the metabolites that can restore growth in each design.

**S3 Data. Mutations**. The breseq identified mutations for all samples collected in this work. Both the full output and a table with only mutations observed in the endpoint clones are provided.

**S4 Data. Duplications**. Genes with read coverage meeting the duplication criteria. Seperate spreadsheets are provided for all samples using the mutant pair, ale number, flask number, isolate number, and replicate number to identify each sample.

## Citations

1. Rittmann bE, Hausner M, Löffler F, Love NG, Muyzer G, Okabe S, et al. A vista for microbial ecology and environmental Biotechnology. Environ Sci Technol. 2006;40: 1096–1103.

2. Minty JJ, Singer ME, Scholz SA, Bae C-H, Ahn J-H, Foster CE, et al. Design and characterization of synthetic fungal-bacterial consortia for direct production of isobutanol from cellulosic biomass. Proc Natl Acad Sci U S A. 2013;110: 14592–14597.

3. Bernstein HC, Carlson RP. Microbial Consortia Engineering for Cellular Factories: in vitro to in silico systems. Comput Struct Biotechnol J. 2012;3: e201210017.

4. Zuroff TR, Xiques Sb, Curtis WR. Consortia-mediated Bioprocessing of cellulose to ethanol with a symbiotic Clostridium phytofermentans/yeast co-culture. biotechnol biofuels. 2013;6: 59.

5. Briones A, Raskin L. Diversity and dynamics of microbial communities in engineered environments and their implications for process stability. Curr Opin Biotechnol. 2003;14: 270–276.

6. Zhang H, Pereira b, Li Z, Stephanopoulos G. Engineering Escherichia coli coculture systems for the production of Biochemical products. Proc Natl Acad Sci U S A. 2015;112: 8266–8271.

7. Zhou K, Qiao K, Edgar S, Stephanopoulos G. Distributing a metabolic pathway among a microbial consortium enhances production of natural products. Nat Biotechnol. 2015;33: 377–383.

8. Saini M, Chen MH, Chiang C-J, Chao Y-P. Potential production platform of n-butanol in Escherichia coli. Metab Eng. 2015;27: 76–82.

9. Flint HJ. The impact of nutrition on the human microbiome. Nutr Rev. 2012;70: S10–S13.

10. Magnúsdóttir S, Heinken A, Kutt L, Ravcheev DA, Bauer E, Noronha A, et al. Generation of genome-scale metabolic reconstructions for 773 members of the human gut microbiota. Nat Biotechnol. 2017;35: 81–89.

11. Adamowicz EM, Hunter RC, Flynn J, Harcombe WR. Cross-feeding modulates antibiotic tolerance in bacterial communities [Internet]. bioRxiv. 2018. p. 243949. doi:10.1101/243949

12. Hosoda K, Suzuki S, Yamauchi Y, Shiroguchi Y, Kashiwagi A, Ono N, et al. Cooperative adaptation to establishment of a synthetic bacterial mutualism. PLoS One. 2011;6: e17105.

13. Hosoda K, Yomo T. Designing symbiosis. Bioeng bugs. 2011;2: 338–341.

14. Mee MT, Collins JJ, Church GM, Wang HH. Syntrophic exchange in synthetic microbial communities. Proc Natl Acad Sci U S A. 2014;111: E2149–56.

15. Baba T, Ara T, Hasegawa M, Takai Y, Okumura Y, Baba M, et al. Construction of Escherichia coli K-12 in-frame, single-gene knockout mutants: the Keio collection. Mol Syst Biol. 2006;2: 2006.0008.

16. Wintermute EH, Silver PA. Emergent cooperation in microbial metabolism. Mol Syst Biol. 2010;6: 407.

17. Zhang X, Reed JL. Adaptive evolution of synthetic cooperating communities improves growth performance. PLoS One. 2014;9: e108297.

18. Hillesland KL, Stahl DA. Rapid evolution of stability and productivity at the origin of a microbial mutualism. Proc Natl Acad Sci U S A. 2010;107: 2124–2129.

19. Zomorrodi AR, Segrè D. Synthetic Ecology of Microbes: Mathematical Models and Applications. J Mol Biol. 2016;428: 837–861.

20. Perez-Garcia O, Lear G, Singhal N. Metabolic Network Modeling of Microbial Interactions in Natural and Engineered Environmental Systems. Front Microbiol. 2016;7: 673.

21. Klitgord N, Segrè D. Environments that induce synthetic microbial ecosystems. PLoS Comput biol. 2010;6: e1001002.

22. Freilich S, Zarecki R, Eilam O, Segal ES, Henry CS, Kupiec M, et al. Competitive and cooperative metabolic interactions in bacterial communities. Nat Commun. 2011;2: 589.

23. Chan SHJ, Simons MN, Maranas CD. SteadyCom: Predicting microbial abundances while ensuring community stability. PLoS Comput Biol. 2017;13: e1005539.

24. Chiu H-C, Levy R, borenstein E. Emergent Biosynthetic capacity in simple microbial communities. PLoS Comput Biol. 2014;10: e1003695.

25. Harcombe WR, Riehl WJ, Dukovski I, Granger bR, betts A, Lang AH, et al. Metabolic resource allocation in individual microbes determines ecosystem interactions and spatial dynamics. Cell Rep. 2014;7: 1104–1115.

26. Zomorrodi AR, Segrè D. Genome-driven evolutionary game theory helps understand the rise of metabolic interdependencies in microbial communities. Nat Commun. 2017;8: 1563.

27. Zomorrodi AR, Maranas CD. OptCom: a multi-level optimization framework for the metabolic modeling and analysis of microbial communities. PLoS Comput biol. 2012;8: e1002363.

28. Feist AM, Palsson bO. The biomass objective function. Curr Opin MicroBiol. 2010;13: 344–349.

29. biliouris K, Babson D, Schmidt-Dannert C, Kaznessis YN. Stochastic simulations of a synthetic bacteria-yeast ecosystem. BMC Syst Biol. 2012;6: 58.

30. Oliveira NM, Niehus R, Foster KR. Evolutionary limits to cooperation in microbial communities. Proc Natl Acad Sci U S A. 2014;111: 17941–17946.

31. Germerodt S, bohl K, Lück A, Pande S, Schröter A, Kaleta C, et al. Pervasive Selection for Cooperative Cross-Feeding in Bacterial Communities. PLoS Comput Biol. 2016;12: e1004986.

32. Lloyd CJ, Ebrahim A, Yang L, King ZA, Catoiu E, O’brien EJ, et al. CObRAme: A Computational Framework for building and Manipulating Models of Metabolism and Gene Expression [Internet]. BioRxiv. 2017. p. 106559. doi:10.1101/106559

33. O{middle dot}brien EJ, middle dot, Lerman JA, Chang RL, Hyduke DR, Palsson bO. Genome-scale models of metabolism and gene expression extend and refine growth phenotype prediction. Mol Syst Biol. 2014;9: 693–693.

34. Lerman JA, Hyduke DR, Latif H, Portnoy VA, Lewis NE, Orth JD, et al. In silico method for modelling metabolism and gene product expression at genome scale. Nat Commun. 2012;3: 929.

35. Wilson M, Lindow SE. Coexistence among Epiphytic bacterial Populations Mediated through Nutritional Resource Partitioning. Appl Environ Microbiol. 1994;60: 4468–4477.

36. Zhao Q, Segre D, Paschalidisy IC. Optimal allocation of metabolic functions among organisms in a microbial ecosystem. 2016 IEEE 55th Conference on Decision and Control (CDC). 2016. doi:10.1109/cdc.2016.7799357

37. Teague bP, Weiss R. SYNTHETIC BIOLOGY. Synthetic communities, the sum of parts. Science. 2015;349: 924–925.

38. Polz MF, Cordero OX. bacterial evolution: Genomics of metabolic trade-offs. Nat Microbiol. 2016;1: 16181.

39. Feist AM, Zielinski DC, Orth JD, Schellenberger J, Herrgard MJ, Palsson bO. Model-driven evaluation of the production potential for growth-coupled products of Escherichia coli. Metab Eng. 2010;12: 173–186.

40. Tepper N, Shlomi T. Predicting metabolic engineering knockout strategies for chemical production: accounting for competing pathways. Bioinformatics. 2010;26: 536–543.

41. Orth JD, Conrad TM, Na J, Lerman JA, Nam H, Feist AM, et al. A comprehensive genome-scale reconstruction of Escherichia coli metabolism––2011. Mol Syst Biol. 2011;7: 535.

42. Zengler K, Zaramela LS. The social network of microorganisms — how auxotrophies shape complex communities. Nat Rev Microbiol. 2018; doi:10.1038/s41579-018-0004-5

43. Fotheringham IG, Dacey SA, Taylor PP, Smith TJ, Hunter MG, Finlay ME, et al. The cloning and sequence analysis of the aspC and tyrB genes from Escherichia coli K12. Comparison of the primary structures of the aspartate aminotransferase and aromatic aminotransferase of E. coli with those of the pig aspartate aminotransferase isoenzymes. Biochem J. 1986;234: 593–604.

44. Thèze J, Margarita D, Cohen GN, Borne F, Patte JC. Mapping of the structural genes of the three aspartokinases and of the two homoserine dehydrogenases of Escherichia coli K-12. J Bacteriol. 1974;117: 133–143.

45. Glansdorff N. Topography Of Cotransducible Arginine Mutations In Escherichia Coli K-12. Genetics. 1965;51: 167–179.

46. Jones-Mortimer MC. Positive control of sulphate reduction in Escherichia coli. Isolation, characterization and mapping oc cysteineless mutants of E. coli K12. Biochem J. 1968;110: 589–595.

47. Sirko AE, Zatyka M, Hulanicka MD. Identification of the Escherichia coli cysM gene encoding O-acetylserine sulphydrylase b by cloning with mini-Mu-lac containing a plasmid replicon. J Gen Microbiol. 1987;133: 2719–2725.

48. Somers JM, Amzallag A, Middleton RB. Genetic fine structure of the leucine operon of Escherichia coli K-12. J Bacteriol. 1973;113: 1268–1272.

49. Wild J, Hennig J, Lobocka M, Walczak W, Ktopotowski T. Identification of the dadX gene coding for the predominant isozyme of alanine racemase in Escherichia coli K12. Mol Gen Genet. 1985;198: 315–322.

50. Lee Y-J, Cho J-Y. Genetic manipulation of a primary metabolic pathway for L-ornithine production in Escherichia coli. Biotechnol Lett. 2006;28: 1849–1856.

51. Felton J, Michaelis S, Wright A. Mutations in two unlinked genes are required to produce asparagine auxotrophy in Escherichia coli. J Bacteriol. 1980;142: 221–228.

52. Vander Horn Pb, backstrom AD, Stewart V, begley TP. Structural genes for thiamine Biosynthetic enzymes (thiCEFGH) in Escherichia coli K-12. J bacteriol. 1993;175: 982–992.

53. Cronan JE Jr, Littel KJ, Jackowski S. Genetic and Biochemical analyses of pantothenate Biosynthesis in Escherichia coli and Salmonella typhimurium. J bacteriol. 1982;149: 916–922.

54. Yang Y, Tsui HC, Man TK, Winkler ME. Identification and function of the pdxY gene, which encodes a novel pyridoxal kinase involved in the salvage pathway of pyridoxal 5’-phosphate Biosynthesis in Escherichia coli K-12. J Bacteriol. 1998;180: 1814–1821.

55. Deatherage DE, barrick JE. Identification of mutations in laboratory-evolved microbes from next-generation sequencing data using breseq. Methods Mol Biol. 2014;1151: 165–188.

56. Ardeshir F, Ames GF. Cloning of the histidine transport genes from Salmonella typhimurium and characterization of an analogous transport system in Escherichia coli. J Supramol Struct. 1980;13: 117–130.

57. Yao N, Trakhanov S, Quiocho FA. Refined 1.89-A structure of the histidine-binding protein complexed with histidine and its relationship with many other active transport/chemosensory proteins. Biochemistry. 1994;33: 4769–4779.

58. Caldara M, Charlier D, Cunin R. The arginine regulon of Escherichia coli: whole-system transcriptome analysis discovers new genes and provides an integrated view of arginine regulation. Microbiology. 2006;152: 3343–3354.

59. Seol W, Shatkin AJ. Escherichia coli alpha-ketoglutarate permease is a constitutively expressed proton symporter. J Biol Chem. 1992;267: 6409–6413.

60. Seol W, Shatkin AJ. Membrane topology model of Escherichia coli alpha-ketoglutarate permease by phoA fusion analysis. J Bacteriol. 1993;175: 565–567.

61. Baker KE, Ditullio KP, Neuhard J, Kelln RA. Utilization of orotate as a pyrimidine source by Salmonella typhimurium and Escherichia coli requires the dicarboxylate transport protein encoded by dctA. J Bacteriol. 1996;178: 7099–7105.

62. The UniProt Consortium. UniProt: the universal protein knowledgebase. Nucleic Acids Res. 2018; doi:10.1093/nar/gky092

63. Riley M, Abe T, Arnaud MB, Berlyn MKB, Blattner FR, Chaudhuri RR, et al. Escherichia coli K-12: a cooperatively developed annotation snapshot--2005. Nucleic Acids Res. 2006;34: 1–9.

64. van Heeswijk WC, Westerhoff HV, Boogerd FC. Nitrogen assimilation in Escherichia coli: putting molecular data into a systems perspective. Microbiol Mol Biol Rev. 2013;77: 628–695.

65. Javelle A, Severi E, Thornton J, Merrick M. Ammonium sensing in Escherichia coli. Role of the ammonium transporter AmtB and AmtB-GlnK complex formation. J Biol Chem. 2004;279: 8530–8538.

66. Finn RD, Coggill P, Eberhardt RY, Eddy SR, Mistry J, Mitchell AL, et al. The Pfam protein families database: towards a more sustainable future. Nucleic Acids Res. 2016;44: D279–85.

67. Song Y, Peisach D, Pioszak AA, Xu Z, Ninfa AJ. Crystal structure of the C-terminal domain of the two-component system transmitter protein nitrogen regulator II (NRII; NtrB), regulator of nitrogen assimilation in Escherichia coli. Biochemistry. 2004;43: 6670–6678.

68. Brown CJ, Todd KM, Rosenzweig RF. Multiple duplications of yeast hexose transport genes in response to selection in a glucose-limited environment. Mol Biol Evol. 1998;15: 931–942.

69. Slack A, Thornton PC, Magner DB, Rosenberg SM, Hastings PJ. On the mechanism of gene amplification induced under stress in Escherichia coli. PLoS Genet. 2006;2: e48.

70. Serres MH, Kerr ARW, McCormack TJ, Riley M. Evolution by leaps: gene duplication in bacteria. Biol Direct. 2009;4: 46.

71. Wallace B, Yang YJ, Hong JS, Lum D. Cloning and sequencing of a gene encoding a glutamate and aspartate carrier of Escherichia coli K-12. J Bacteriol. 1990;172: 3214–3220.

72. Carter EL, Jager L, Gardner L, Hall CC, Willis S, Green JM. Escherichia coli abg genes enable uptake and cleavage of the folate catabolite p-aminobenzoyl-glutamate. J Bacteriol. 2007;189: 3329–3334.

73. Bailey SF, Rodrigue N, Kassen R. The effect of selection environment on the probability of parallel evolution. Mol Biol Evol. 2015;32: 1436–1448.

74. Goldford JE, Lu N, Bajic D, Estrela S, Tikhonov M, Sanchez-Gorostiaga A, et al. Emergent Simplicity in Microbial Community Assembly [Internet]. 2017. doi:10.1101/205831

75. Ebrahim A, Brunk E, Tan J, O’Brien EJ, Kim D, Szubin R, et al. Multi-omic data integration enables discovery of hidden biological regularities. Nat Commun. 2016;7: 13091.

76. Kallus Y, Miller JH, Libby E. Paradoxes in leaky microbial trade. Nat Commun. Nature Publishing Group; 2017;8: 1361.

77. Sheth RU, Cabral V, Chen SP, Wang HH. Manipulating Bacterial Communities by in situ Microbiome Engineering. Trends Genet. 2016;32: 189–200.

78. Mueller UG, Sachs JL. Engineering Microbiomes to Improve Plant and Animal Health. Trends Microbiol. 2015;23: 606–617.

79. Ebrahim A, Lerman JA, Palsson BO, Hyduke DR. COBRApy: COnstraints-Based Reconstruction and Analysis for Python. BMC Syst Biol. 2013;7: 74.

80. Burgard AP, Pharkya P, Maranas CD. Optknock: A bilevel programming framework for identifying gene knockout strategies for microbial strain optimization. Biotechnol Bioeng. 2003;84: 647–657.

81. Orth JD, Palsson B. Gap-filling analysis of the iJO1366 Escherichia coli metabolic network reconstruction for discovery of metabolic functions. BMC Syst Biol. 2012;6: 30.

82. Chen S, Huang T, Zhou Y, Han Y, Xu M, Gu J. AfterQC: automatic filtering, trimming, error removing and quality control for fastq data. BMC Bioinformatics. 2017;18: 80.

83. Grenier F, Matteau D, Baby V, Rodrigue S. Complete Genome Sequence of Escherichia coli BW25113. Genome Announc. 2014;2. doi:10.1128/genomeA.01038-14

84. Langmead B, Salzberg SL. Fast gapped-read alignment with Bowtie 2. Nat Methods. 2012;9: 357–359.

85. Li H, Handsaker B, Wysoker A, Fennell T, Ruan J, Homer N, et al. The Sequence Alignment/Map format and SAMtools. Bioinformatics. 2009;25: 2078–2079.

86. Seabold S, Perktold J. Statsmodels: Econometric and statistical modeling with python. Proceedings of the 9th Python in Science Conference. SciPy society Austin; 2010. p. 61.

87. Yang L, Ma D, Ebrahim A, Lloyd CJ, Saunders MA, Palsson BO. solveME: fast and reliable solution of nonlinear ME models. BMC Bioinformatics. bmcbioinformatics. biomedcentral. …; 2016;17: 391.

88. Ma D, Yang L, Fleming RMT, Thiele I, Palsson BO, Saunders MA. Reliable and efficient solution of genome-scale models of Metabolism and macromolecular Expression. Sci Rep. 2017;7: 40863.

